# Antibiotic transport kinetics in Gram-negative bacteria revealed via single-cell uptake analysis and mathematical modelling

**DOI:** 10.1101/645507

**Authors:** Jehangir Cama, Margaritis Voliotis, Jeremy Metz, Ashley Smith, Jari Iannucci, Ulrich F. Keyser, Krasimira Tsaneva-Atanasova, Stefano Pagliara

## Abstract

The double-membrane cell envelope of Gram-negative bacteria is a formidable barrier to intracellular antibiotic accumulation. A quantitative understanding of antibiotic transport in these cells is crucial for drug development, but this has proved elusive due to the complexity of the problem and a dearth of suitable investigative techniques. Here we combine microfluidics and time-lapse auto-fluorescence microscopy to quantify antibiotic uptake label-free in hundreds of individual *Escherichia coli* cells. By manipulating the microenvironment, we showed that drug (ofloxacin) accumulation is higher in growing versus non-growing cells. Using genetic knockouts, we provide the first direct evidence that growth phase is more important for drug accumulation than the presence or absence of individual transport pathways. We use our experimental results to inform a mathematical model that predicts drug accumulation kinetics in subcellular compartments. These novel experimental and theoretical results pave the way for the rational design of new Gram-negative antibiotics.

## Introduction

Life depends on the exchange of molecules between cells and their surroundings^1^. Cells have evolved elaborate, adaptable envelope structures to optimize nutrient accumulation while restricting the uptake of xenobiotics, particularly those that negatively impact their survival. However, it is these very attributes that make the study of these molecular transport processes extremely challenging. Transport across the cell envelope may occur passively via diffusion^2^, either through lipids or specific protein pores^3^, or via active transporters^4^, which move substrates both into and out of the cell. Furthermore, the expression of these different pathways is often strongly regulated by the surrounding microenvironment^5^ and can vary from cell to cell^6^. Due to the many complexities of studying these transport problems, biophysical and mathematical modelling has been used extensively to uncover detailed features of molecular transport in synthetic model systems. For instance, a mathematical study of hydrodynamic entrance effects showed that the hourglass shape of aquaporins might be a result of natural selection processes optimizing water permeability^7^. One-dimensional diffusional models, both theoretical^8^ and experimental^9^ have been used to shed light on the single-file motion of particles through narrow constrictions, simulating molecular transport through biological nanopores. Colloidal model systems have been used to investigate Brownian dynamics in biomimetic systems^10^, with recent reports showing the breakdown of transition-path-time symmetry on molecular and meso-scales out of equilibrium^11^.

However, these molecular-scale modelling studies do not capture the kinetics of substrate uptake in living cells and, from a biomedical perspective, a key transport challenge involves quantitatively understanding the intracellular uptake of antibiotics in bacteria^12,13^. Antibiotic failure in the treatment of microbial infections is predicted to cause 10 million deaths *annually* by 2050^14^. Gram-negative bacterial infections are of particular concern, due to the protection against antibiotics provided by their complex double-membrane cell envelopes (Figure 1A). These structures include an asymmetric outer membrane that contains lipopolysaccharide (LPS) molecules, which create a formidable permeability barrier to the cellular entry of both hydrophilic and hydrophobic molecules^12,15^. Antibiotic permeation across the outer membrane is therefore dependent on the drug’s ability to utilize protein pores (or *porins*)^3,16,17^, typically used for nutrient uptake, to circumvent this barrier. These porins show a preference for hydrophilic, charged compounds; however, antibiotics that are active against targets located in the cytoplasm have to also cross the inner membrane phospholipid bilayer, which acts as a selectivity barrier *against* polar, charged molecules^12,15^. Additionally, Gram-negative bacteria harbor active efflux mechanisms, which pump toxic compounds out of the cell^18^. Successful drugs must minimize their propensity for recognition and removal by these efflux pumps, in addition to displaying specific physicochemical properties to permeate both through the outer membrane porins and inner membrane phospholipids^15^.

**Fig. 1.**
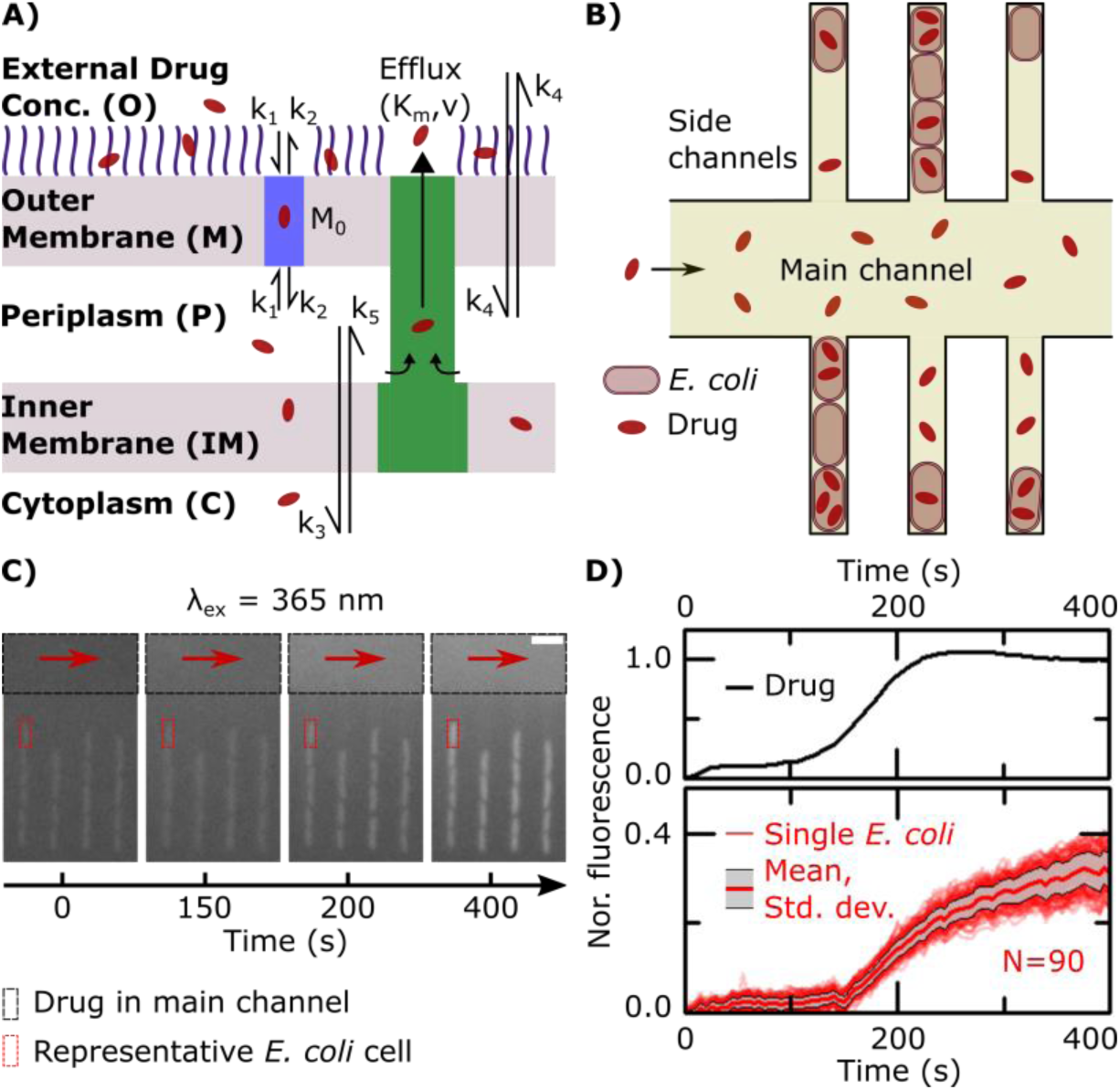
Quantifying and modelling ofloxacin uptake label-free in individual *E. coli* cells. **A)** Schematic of the main processes involved in drug translocation across Gram-negative cell envelopes. Drug molecules penetrate the outer membrane (M) primarily through protein porins, with association and dissociation rates *k*_1_ and *k*_2_, respectively. *M*_0_ refers to the concentration of functional porin binding sites in the outer membrane. Any residual (non-porin) transport across the outer membrane LPS barrier is modelled with *k*_4_. Drug transport through the inner membrane is modelled with kinetic parameters *k*_3_ and *k*_5_. Drug molecules are subject to removal from the cell via active efflux mechanisms which follow Michaelis-Menten kinetics (*K*_*m*_, *v*). **B)** Schematic of the microfluidic chip used for the ofloxacin uptake experiment. A main channel of height 25 μm and width 100 μm is used for continuously exchanging the microenvironment with nutrient, drug or dye delivery; cells are confined single-file in a network of side channels whose height and width are both 1.4 μm, with length 25 μm. **C)** Section of epifluorescence images showing the delivery of ofloxacin (100×MIC, 12.5 μg/ml in PBS) and its corresponding uptake by the cells in the side channels. The ofloxacin molecules within and around the bacteria are tracked using their auto-fluorescence at λ_ex_= 365 nm. Scale bar = 5 μm. **D)** Quantitative estimation of the temporal profile of ofloxacin delivery in the chip, and the corresponding ofloxacin uptake profile of 90 individual *E. coli* cells; the thick red line represents the mean and the grey shaded area the standard deviation of the ofloxacin uptake profiles of the 90 cells investigated. The fluorescence values are reported after correcting for the background and normalizing to the fluorescence of the drug as detailed in the Methods. The complete datasets prior to normalization for the three different *E. coli* strains investigated are presented in the SI in Figure S6.

The study of drug uptake is further complicated by the fact that the expression and activity of porins and efflux pumps vary i) with the microenvironment conditions^5^ and ii) within an isogenic population exposed to the same environmental landscape^19^. Many existing experimental techniques suffer from the requirement of complex washing steps^12,13^, with cells only studied after resuspension in contrived nutrient environments^20,21^; the washes also increase the chance of cell lysis and efflux or diffusion of the analyte from the cells, besides affecting cellular physiology. Furthermore, the most commonly used techniques are population level assays which cannot investigate uptake at the single-cell or at the subcellular level. Finally, most of the available techniques only provide a static picture of drug accumulation rather than the dynamic evolution of drug uptake. There is therefore a need to fundamentally change the experimental approach for quantifying antibiotic accumulation in individual bacteria after exposure to different nutrient conditions or in different metabolic states. Ideally, this approach should also be simple to implement to ensure its uptake in pharmaceutical companies and in clinical settings.

Here, we address these myriad challenges by introducing a unique combination of single-cell uptake analysis and mathematical modelling to study drug accumulation and kinetics in up to hundreds of individual cells per experiment. To do so we used *Escherichia coli* as a model organism for Gram-negative bacteria, seeded a small aliquot of bacterial culture into a microfluidic “mother-machine” device^22^ (Figure 1B) and dosed *E. coli* either in a non-growing or a growing state with the fluoroquinolone antibiotic ofloxacin (12.5 μg/ml) while imaging the kinetics of ofloxacin accumulation in individual *E. coli* (Figure 1C-D) using the auto-fluorescence of the drug. Quinolones such as ofloxacin disrupt the DNA replication process in the cytoplasm of bacteria; in *E. coli*, the primary target is the enzyme DNA gyrase, a tetramer which is composed of two copies each of its subunits, GyrA and GyrB^23^. Therefore ofloxacin activity depends directly on its ability to accumulate in the cytoplasm.

Using biophysical experimental model systems, we and others have previously shown *in vitro* that porins such as OmpF facilitate quinolone transport across the outer membrane^16,24^, and that quinolones also diffuse freely across phospholipid bilayers such as those found in the cytoplasmic membrane^25^. However, the role of the TolC efflux protein in quinolone transport is currently a matter of debate. Although a *tolC* deficient strain of a fluoroquinolone-resistant clinical *E. coli* isolate was shown to be more susceptible to fluoroquinolones than the parental strain^26^, TolC levels alone do not necessarily limit drug efflux capabilities in *E. coli*^27^. Cellular quinolone accumulation data comparing parental strains and their corresponding *tolC* knockouts also show contradictions, with some reports showing increased accumulation^26^ in the knockout and others showing no significant differences between the strains^28^.

We use our novel approach to investigate this complex membrane transport landscape by performing ofloxacin accumulation experiments in three *E. coli* strains from the Keio collection^29^, encompassing the parental strain (PS) BW25113, an OmpF porin knockout (Δ*ompF*) and a TolC efflux protein knockout (Δ*tolC*) strain. We confirmed that OmpF plays a significant role in ofloxacin transport^30^, but found that the absence of TolC appears to have no significant impact on drug accumulation compared to the PS. Even more surprisingly, our ability to directly compare the role of these transport proteins and the nutrient environment in drug uptake revealed, for the first time, that the microenvironment affects ofloxacin accumulation to a greater extent than the loss of the key transport pathways that we investigated.

Furthermore we applied a set of three ordinary differential equations to model the uptake process^31^ across the three strains in order to complement our experiments. This allowed us to estimate the kinetic parameters associated with early stage ofloxacin uptake. We combined this with Bayesian inference to investigate how specific model parameters varied between individual cells in the different strains. We used the parameters obtained from the modelling and statistical inference to *predict* the kinetics of drug accumulation in the various subcellular compartments of the cells across the different strains. For the avoidance of any confusion, we stress that the modelling results are *theoretical* results^32^ *inferred* from our experimental data which provide *predictions* of the levels of subcellular drug accumulation; the experimental validation of these predictions is beyond the scope of any currently available technology, particularly at the single-cell level. Finally, although this study focuses on Gram-negative bacteria, the experimental and theoretical framework that we employ may be repurposed, with appropriate modifications, for advancing our understanding of molecular transport in a range of fundamental phenomena in both cellular and synthetic systems. This will pave the way for a direct, quantitative evaluation of the role of growth phases, nutrient conditions and transport pathways in drug accumulation in cells.

## Results

Figure 2A-D report bacterial drug uptake profiles (red lines) from representative experiments studying growing PS (2A), non-growing PS (2B), growing Δ*ompF* (2C) and growing Δ*tolC* (2D) *E. coli*. The drug uptake profiles for Δ*tolC* (non-growing) *E. coli* and all the biological repeats performed are reported in Figure S6. We quantify drug dosage precisely via its fluorescence (SI Note 1) in every experiment. Further, we performed cellular autofluorescence controls in the absence of the drug and show that this has a negligible effect on our results (SI Note 2).

**Fig. 2.**
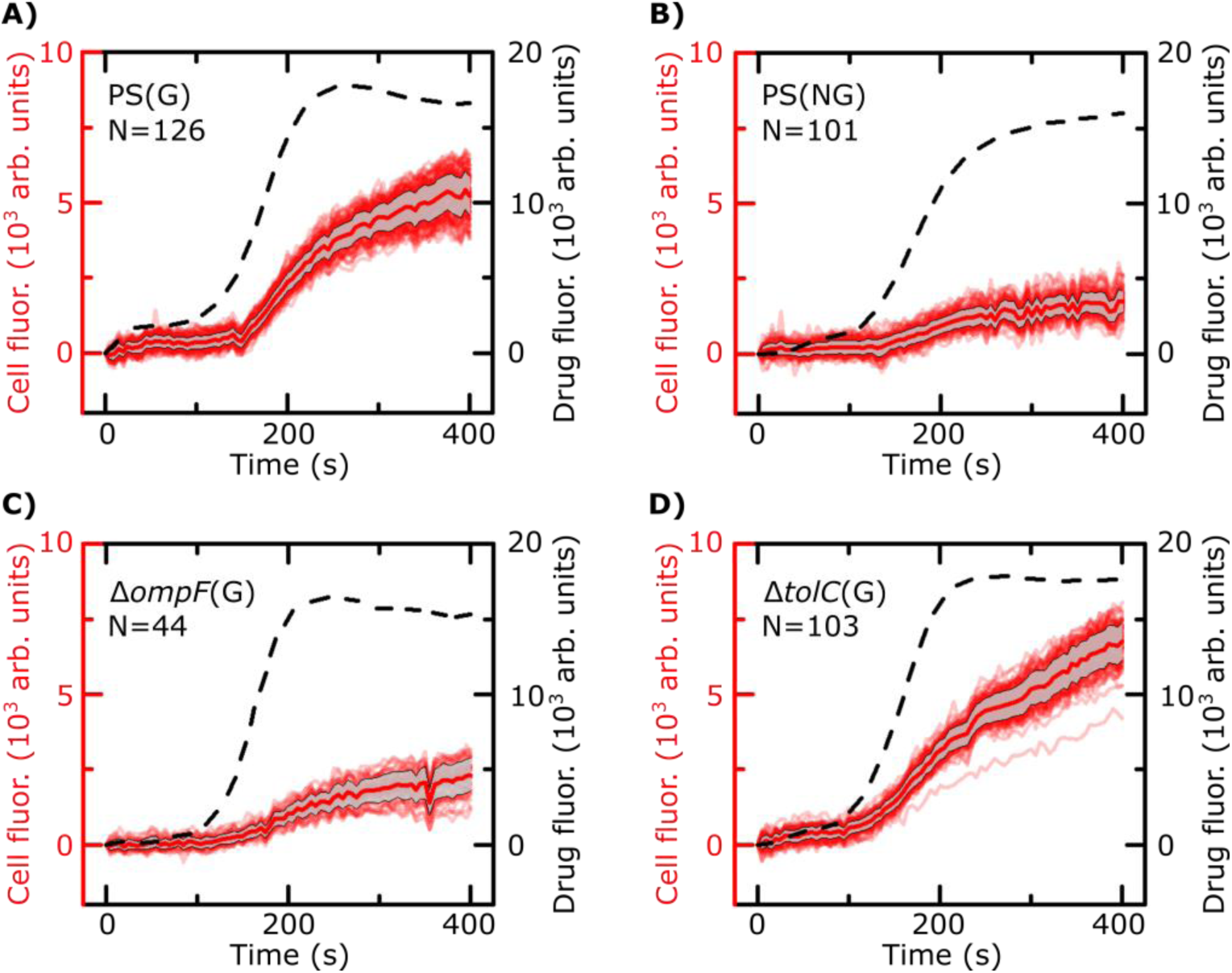
Representative ofloxacin uptake experiments for the bacterial strains/conditions investigated. The *E. coli* strains (Parental Strain, PS; Δ*ompF*; Δ*tolC*), conditions (growing, G; non-growing, NG) and number of cells (*N*) are indicated inset. All values are reported after subtracting the background and the initial cellular fluorescence (before drug arrival) as explained in the Methods. For reference, the complete datasets for all strains/conditions including all the biological repeats are provided in Figure S6 in the SI. Dashed lines represent the drug dosage profiles (right Y-axes) in the main channel. These individual drug dosage profiles are provided as inputs when modelling the drug uptake in the corresponding cells in an experiment. The cell fluorescence profiles are shown in red (left Y-axes), along with the mean (thick red line) and standard deviation (grey shading) for all the cells in an experiment. Comparing growing versus non-growing PS bacteria (panels A and B) directly shows that the growing cells accumulate more drug than non-growing cells. This is apparent in the Δ*tolC* strain as well (Figure S6). Comparing the cell fluorescence profiles of growing PS (A), Δ*ompF* (C) and Δ*tolC* (D) also clearly shows that the Δ*ompF* mutant accumulates less ofloxacin than the other two strains. A quantitative analysis of the amount of drug accumulated at the end of the experiments for each strain/condition is provided in Figure 3.

We observe an increase in cellular drug fluorescence within seconds after the arrival of the drug in the vicinity of the cells. Please note that previous population-level studies have shown biphasic ofloxacin uptake in *E. coli* over longer timescales of up to an hour^33^, but here we focus our attention on the initial stages of drug uptake, studying the immediate cellular response to drug dosage (*t* ≤ 400 s) at the single-cell level.

### 1. Growing bacteria accumulate more ofloxacin than non-growing bacteria

Comparing growing versus non-growing PS cells (Figure 2A-B) immediately reveals that growing cells accumulate more ofloxacin than non-growing cells. To quantify this difference, we compared the distributions of cellular fluorescence (normalized to the value of drug fluorescence) at *t* = 400 s across all experimental repeats in Figure 3 (see Methods). In all datasets, growing PS cells show an approximately 3-fold higher fluorescence than non-growing cells (**growing**: norm. fluor. = 0.34 ± 0.11, *N* = 317, mean ± s.d.; **non-growing**: norm. fluor. = 0.10 ± 0.03, mean ± s.d., *N* = 405; p<10^−10^). A similar result was obtained when comparing growing and non-growing cells in the Δ*tolC* mutant strain (**growing**: norm. fluor. = 0.31 ± 0.08, *N* = 211, mean ± s.d.; **non-growing**: norm. fluor. = 0.12 ± 0.06, mean ± s.d., *N* = 193; p<10^−10^).

**Fig. 3.**
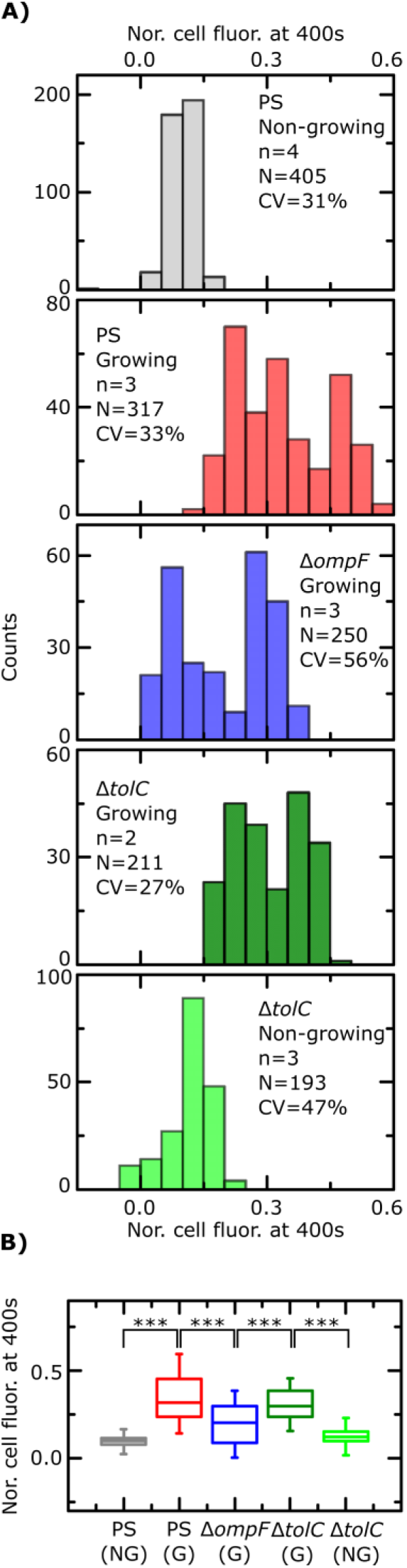
Final level of normalized whole cell fluorescence for the different strains and nutritional conditions. **(A)** Fluorescence distributions across the different strains and conditions. In the insets, *n* refers to number of experimental repeats, *N* reports the total number of bacteria and *CV* refers to the coefficient of variation of the data. All comparisons are made at *t* = 400 s. **(B)** Comparison of data pooled from the different experiments shows that non-growing PS *E. coli* show significantly lower ofloxacin uptake than growing PS *E. coli* (*p*<10^−10^). This was also true in the Δ*tolC* strain, where non-growing cells showed significantly lower uptake (*p*<10^−10^) than growing cells, suggesting ofloxacin uptake critically depends on the growth phase of the cells within the timescales of our experiment. Growing Δ*ompF E. coli* showed lower whole cell drug accumulation than growing PS (*p*<10^−10^) and Δ*tolC* (*p*<10^−10^) cells, in line with expectations. However, growing Δ*ompF E. coli* accumulated *more* ofloxacin than non-growing PS cells (*p*<10^−10^), suggesting that the growth phase of the cells as set by the nutrient environment plays an even more important role than the deletion of *ompF* in drug uptake. The horizontal lines in the interior of the boxes report the medians of the respective distributions. Statistical significance tested using a 2-sample t-test incorporating Welch’s correction; the complete set of p-values is reported in the SI (Table S1).

### 2. Knocking out *ompF* lowers ofloxacin accumulation compared to the PS

From Figure 2A and 2C, we also observe that the growing Δ*ompF* mutant strain accumulates lower amounts of ofloxacin than the PS (growing) over the timescales investigated. This is quantified in Figure 3 (**Δ*ompF***: norm. fluor. = 0.20 ± 0.11, mean ± s.d., *N* = 250; **PS**: norm. fluor. = 0.34 ± 0.11, *N* = 317, mean ± s.d.; p<10^−10^); knocking out the OmpF porin thus lowers the ability of ofloxacin to permeate into the cell compared to the parental strain. Our result agrees with previous reports that show that OmpF facilitates fluoroquinolone transport across Gram-negative outer membranes^3,24^.

### 3. Knocking out *tolC* does *not* increase ofloxacin accumulation compared to the PS

Interestingly, we were unable to detect an increase in ofloxacin accumulation in growing Δ*tolC* mutant cells compared to the PS at the 400 s time-point (Figure 3). In fact, as reported above, we measured a small *decrease* in the drug fluorescence in growing Δ*tolC* cells compared to the growing PS cells (**Δ*tolC***: norm. fluor. = 0.31 ± 0.08, *N* = 211, mean ± s.d.; **PS**: norm. fluor. = 0.34 ± 0.11, *N* = 317, mean ± s.d.; p=2.7×10^−4^). This finding is addressed in detail in the Discussion.

### 4. Direct comparison reveals that growth phase plays a more significant role in ofloxacin accumulation than knocking out *ompF*

Our ability to directly compare drug accumulation in different metabolic states revealed that the growing Δ*ompF* mutant strain accumulates more ofloxacin than the non-growing PS (**growing Δ*ompF***: norm. fluor. = 0.20 ± 0.11, mean ± s.d., *N* = 250; **non-growing PS**: norm. fluor. = 0.10 ± 0.03, *N* = 405, mean ± s.d.; p<10^−10^), suggesting that the growth phase plays an even bigger role than the removal of OmpF in drug uptake. We believe this is the first time such a direct comparison has been performed. These results emphasize the importance of studying the role of the cellular metabolic state in drug uptake.

### 5. Ofloxacin uptake is homogeneous across a clonal population

A major advantage of single-cell approaches is their ability to quantify heterogeneity (or the lack thereof) in the cellular response to treatment within the individual cells in a population^34^. In order to estimate heterogeneity in drug uptake across the bacteria, we first estimated the variation in cellular fluorescence in the absence of the drug and found a mean coefficient of variation (CV) of approximately 10% (see Methods). We found a similar CV when quantifying the heterogeneity in the cellular fluorescence corresponding to drug uptake. As seen in Figure S6, such variation is representative across the biological repeats. We thus conclude that ofloxacin uptake is homogeneous across the clonal populations that we studied, which is remarkable considering the recent reports on cellular heterogeneity within microbial populations^19^, including considerable heterogeneity in glucose uptake in *E. coli* cells^35^.

### Theoretical predictions from a mathematical model of drug transport across the Gram-negative cell envelope

The quantitative comparisons above provide a *static* picture regarding the impact of porins, pumps and growth stages on ofloxacin accumulation in Gram-negative bacteria at the *whole-cell level*. However, the most desirable information concerns the dynamics governed by the *kinetics* of drug accumulation in different *subcellular compartments*. It is crucial to understand how much of a drug actually reaches its target which, in the case of ofloxacin, lies in the cytoplasm^36^. However, there are currently no experimental techniques capable of quantifying subcellular drug accumulation at the single-cell level. We therefore turn to theoretical modelling to investigate this process. We rationalize our experimental single-cell drug uptake data via a mathematical model (see Methods), where parameters governing porins (*M*_0_) and efflux pumps (*v*) are allowed to vary between cells in the population according to a log-normal distribution^37^. The inferred parameter distributions for growing bacteria from the three investigated strains are presented in Figure 4A-B; the different experimental repeats are signified by solid, dotted and dashed lines (PS, red; Δ*ompF*, blue; Δ*tolC*, green). We found similar values across the different replicates for the PS cells, whereas the knockout mutants showed greater variability both between replicates and within individual experiments, as observed in Figure 4A-B. The parameter estimations also confirmed lower porin concentrations in the Δ*ompF* mutant compared to the PS. Note that due to the flatness of the uptake profiles of the non-growing cells, we chose not to infer model parameters from those experiments.

**Fig. 4.**
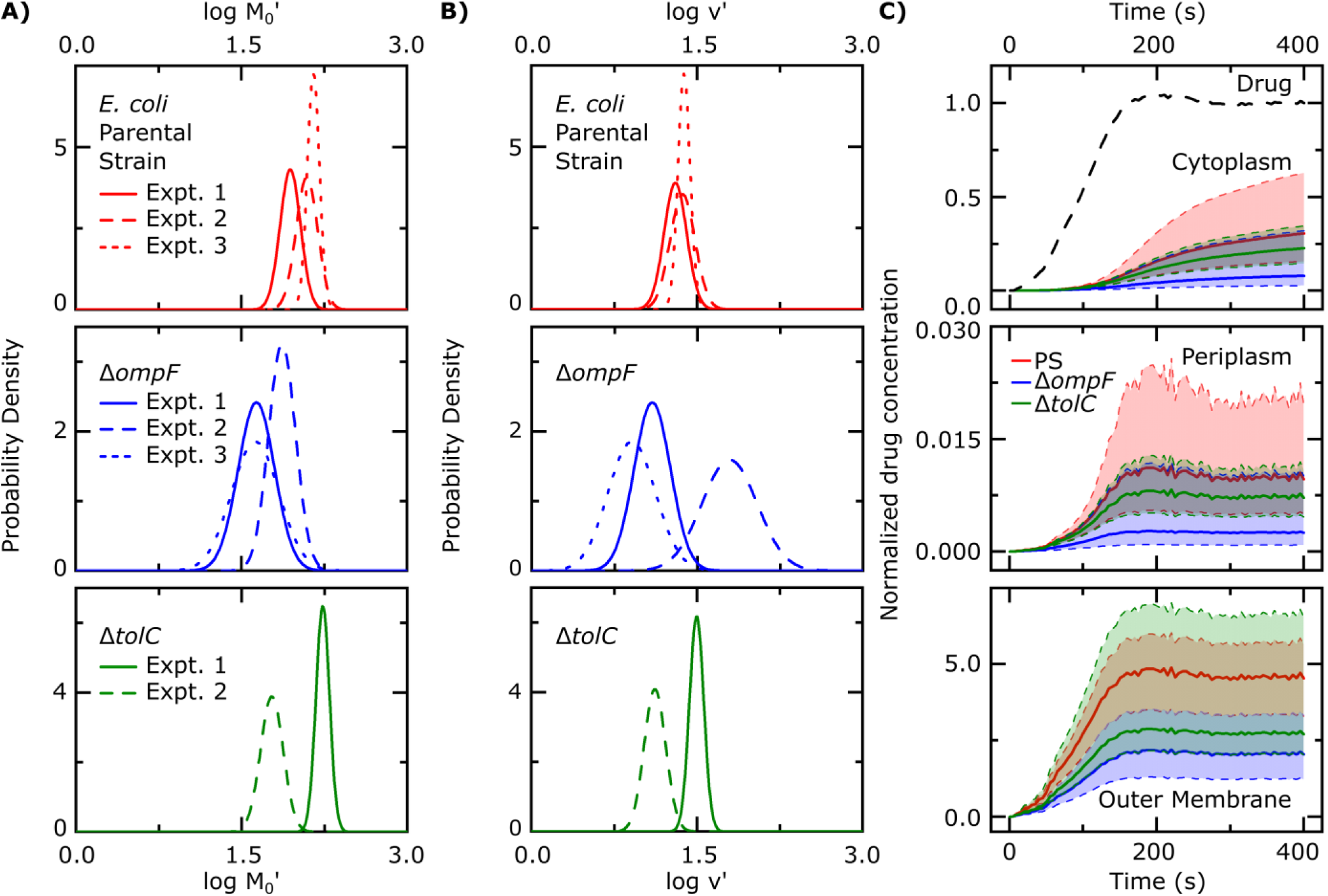
Drug accumulation kinetics predicted by fitting the single-cell data to the drug uptake model. Maximum aposteriori estimates of the population distribution of parameters 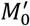**(A)** and *v*′ **(B)** for growing parental strain (PS, top), Δ*ompF* (middle) and Δ*tolC* (bottom) *E. coli*. The solid, dashed and dotted lines refer to individual experimental repeats. These distributions were generated using the mode of the joint posterior distribution of the means and standard deviations of the log-normal distributions for 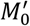 and *v*′; the marginal posterior distributions of the means and standard deviations for the parameters are provided in the SI in Figures S4 and S5 respectively. **C)** Predicted ofloxacin uptake in the different bacterial compartments. Temporal dependence of the normalized drug concentration in the cytoplasm, periplasm and outer membrane for PS (red), Δ*ompF* (blue) and Δ*tolC* (green) bacteria in response to the drug dosage input (dashed black line, top panel). These drug uptake profiles were obtained by using the kinetic parameter values in (A) and (B) and the theoretical model (equations (i)-(iii)). The concentrations reported are normalized to the drug dosage concentration (12.5 μg/ml ofloxacin). The solid lines correspond to median accumulation in the respective compartments and the shaded area represents the [20,80] posterior predictive interval of the accumulation. The results shown were generated by running the model using 500 independent samples of parameters 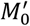 and *v*′ from their joint posterior distributions. All other parameters were fixed to the values given in Table S2. The model predicts the saturation of binding sites in the outer membrane. The median saturation concentration in the outer membrane is approximately 2.25-fold higher in the PS compared with the Δ*ompF* strain. The periplasmic drug concentrations are approximately 30-fold lower than the cytoplasmic concentrations, which is likely due to the drug binding to its targets within the cytoplasm. Using the [20,80] posterior predictive intervals, we have calculated the probabilities of cells from the different strains showing higher/lower accumulation in the different compartments in a pairwise manner. These results are provided in Table S4.

Once model parameters were inferred from all the individual experiments (using the corresponding drug dosage profiles for each experiment), we used these parameters in the model to predict drug accumulation in the various subcellular compartments for cells belonging to the three strains (Figure 4C). In this estimation for Figure 4C, we used an average experimental drug dosage profile (dashed black line, top panel, Figure 4C) as the input. The overlap (or lack thereof) between the [20,80] posterior predictive intervals (shaded regions in Figure 4C) allows us to predict the probability of PS cells having a higher/lower ofloxacin concentration than each of the mutants, at the subcellular level. The pairwise comparisons (at *t* = 400 s) for the different strains/compartments are presented in Table S4.

The model predicts that the drug saturates all the binding sites in the outer membrane within approximately 175 s in all three strains. The PS strain has the highest outer membrane drug concentration, with the Δ*ompF* mutant having an approximately 2.25-fold lower concentration, which corresponds to the fewer binding sites available in the mutant (Figure 4A). At the end of the experiment, the probability that the PS strain has a higher drug concentration than the Δ*ompF* mutant in the outer membrane is 0.924; in contrast, between the PS and the Δ*tolC* mutant, the probability that the PS has more drug in the outer membrane is 0.525, suggesting no appreciable difference (Table S4).

The periplasm is also predicted to contain approximately 30-fold lower ofloxacin concentrations than the cytoplasm for all three strains at *t* = 400 s – this is likely due to the binding of the ofloxacin molecules to their targets within the cytoplasm. The model also predicts a lag time of approximately 100 s between drug accumulation in the outer membrane versus drug uptake in the cytoplasm. In the cytoplasm, the difference between the PS and the mutant strains is less obvious. The model predicts that, at the end of the experiment, the PS strain has a probability of 0.719 of having a higher drug concentration in the cytoplasm than the Δ*ompF* mutant (Table S4). Comparing the PS and the Δ*tolC* mutant, the corresponding probability is 0.549.

## Discussion

Drug uptake in Gram-negative bacteria is an extremely complex biophysical phenomenon because of the different physicochemical pathways and combination of active and passive transport processes involved. However, it is essential to understand the roles of these pathways in a quantitative manner to rationally design drugs that can accumulate in the vicinity of their targets, which will crucially contribute to overcoming the void in Gram-negative drug discovery.

We have developed a novel combination of experiment and theoretical modelling to tackle the challenge of quantifying antibiotic uptake in single Gram-negative bacteria. Unlike the majority of techniques, which involve complex washing steps after drug delivery, or are limited to certain specific media conditions^12,13^, our microfluidic platform facilitates the study of drug uptake in different microenvironments and cellular metabolic states. We quantify drug dosage in every experiment, which allows us to correct for any variations in fluorescence intensities/flow conditions between experiments. Since we use microfluidics, we quantify drug uptake from the moment the drug arrives in the vicinity of the cells, facilitating the real-time measurement of the transport process.

It is worth noting that we can measure over a hundred cells in an experiment; by reducing the time resolution it is also possible to correspondingly increase the number of cells measured, since typically thousands of cells are confined in the microfluidic device. This ability will be used in future studies, especially for drugs whose uptake timescales are longer than fluoroquinolones. Since our excitation wavelength is 365 nm, in contrast to previous studies using deep UV illumination to study antibiotic uptake in single cells^20,38^, we can work with standard optics and light sources, rather than needing quartz objectives and cover slips, and deep UV light sources which may not be easily accessible. Although cellular metabolites may also fluoresce at similar wavelengths, we have corrected this by subtracting the baseline cellular fluorescence as described in the Methods (and in SI Note 2). Note that metabolite concentrations are known to fluctuate in response to fluoroquinolone treatment, but this is typically less than a two-fold change within the timescales of our experiment and includes both increases and decreases^39^. The baseline cellular autofluorescence (growing PS cells, Figure S1B) shows typical intensities of approximately 1700 (arb. units), while the fluorescence *increases* in the cells due to drug accumulation are approximately 5200 (arb. units, Figure S1C). Therefore, we estimate that the *maximum* contribution of metabolites to our fluorescence signal, in the case where *all* the metabolites were to double in number (and assuming that the fluorescence scales linearly), would be approximately 33% in this case; however, considering that the metabolites show both increases and decreases in response to fluoroquinolone treatment, we estimate that the actual contribution is significantly lower, and would constitute a higher order correction to our results. Note that a non-fluorescent version of ofloxacin does not exist, making a direct measurement of the changes in metabolite autofluorescence in response to ofloxacin treatment intractable. However, we reiterate that the baseline cellular autofluorescence is already accounted for in our analysis.

Using our novel approach, we established that within the timescales investigated, ofloxacin accumulates to a greater degree in growing versus non-growing bacteria (Figures 2 and 3). It is likely that this reduction in ofloxacin accumulation contributes to the significant increase in cell survival to this drug that was previously observed as the cells enter stationary phase compared with early exponential phase cultures^40,41^. In previous work, we profiled the entire transcriptome of *E. coli* (BW25113) growing in LB media at various time points across the growth cycle; this revealed that the expression of the genes encoding the ofloxacin target DNA gyrase (specifically, its subunits GyrA and GyrB) does not change substantially across the growth cycle^40^. This agrees with a previous study which showed that the levels of the Gyr proteins do not change appreciably as cells grow from exponential into stationary phase; indeed, the authors found no appreciable degradation of the Gyr proteins even after 72 h of starvation^42^. However, our transcriptomics revealed that the expression of the genes encoding the major *E. coli* porins OmpF and LamB, through which antibiotics diffuse, was significantly upregulated in exponentially growing compared to stationary phase *E. coli* cells^40^. For convenience, we have reproduced the transcriptomic data of the genes relevant to our study in Figure S7 in the SI. This strongly suggests that the differences in ofloxacin uptake that we observe between growing and non-growing cells are due to phenotypic modifications of the cell envelope transport pathways, rather than phenotypic modifications at the drug target level.

In growing cells, knocking out the *ompF* gene led to a decrease in drug accumulation compared to the parental strain, in line with previous results^3^, confirming that fluoroquinolones utilize porins to enter *E. coli* cells. The model predicts an approximately 4-fold lower median cytoplasmic concentration of ofloxacin in the Δ*ompF* mutant compared to the PS (growing cells) at the end of the experiment (Figure 4C). However, the effect of the growth phase was more significant than the removal of the porin – non-growing PS cells accumulated lower amounts of ofloxacin than the growing Δ*ompF* mutant (Figure 3). Previous studies have reported that nutrient-starved bacteria show reduced drug uptake^43^, but these studies did not determine the extent to which environmental factors, and subsequent cell phenotypic acclimation, predetermine drug uptake compared to genotypic changes which result in protein loss.

As described in the Results, we did not measure any increase in drug accumulation in the Δ*tolC* strain. This is a matter of debate in the literature; as noted in the introduction, different groups have investigated the role of TolC in fluoroquinolone accumulation in *E. coli*, and have reported contradictory results^26,28^. The TolC outer membrane efflux protein forms an important part of multi-drug efflux systems such as AcrAB-TolC that eject antibiotics and other toxins from *E. coli* cells^27^, and naively one would have expected that losing TolC negatively affects the ability of the cell to efflux the antibiotic, thus increasing its intracellular accumulation. It has also been reported that the inactivation of *tolC* increases the susceptibility of bacteria to a range of antibacterial agents, ostensibly due to the inactivation of the corresponding efflux systems^27^. However, although the overproduction of the AcrAB-TolC efflux system has been implicated in the antibiotic resistance of clinical isolates of *E. coli* species, there was no significant correlation between the overexpression of the *acrAB* and *tolC* genes^27,44^. With regards to fluoroquinolone antibiotics, it was reported that average *tolC* expression levels in fluoroquinolone-susceptible and fluoroquinolone-resistant clinical isolates of *E. coli* were not statistically different^27,44^. Zgurskaya and co-workers therefore concluded that TolC quantities alone do not limit the drug efflux capabilities of *E. coli*^27^. Our data further corroborate this hypothesis.

The use of mathematical modelling and Bayesian inference to rationalize our data enabled us to maximize the information embedded in our time-lapse single-cell measurements, leading to predictions of the *kinetics* of the uptake process. We extracted kinetic parameters corresponding to the single-cell drug uptake profiles and quantified changes in these parameters in the different strains (Figure 4A-B). To validate our inference procedure, we used data simulated by the model and showed that we can indeed recover the parameter values which were used for generating these (Fig. S8). Importantly, the model allowed us to predict drug accumulation in the different subcellular compartments, which is a major milestone for the entire research community working on this problem. It is important to note that these are *predictions*, arising out of our whole-cell data; validation of the model predictions regarding subcellular levels of drug concentration will only be possible once the considerable experimental challenges for these measurements at the single-cell level are overcome. There are currently no techniques capable of resolving the concentrations of drugs in different subcellular compartments in individual cells. Future work will also involve studying drug accumulation after modulation of other transport pathways in the Gram-negative double membrane to estimate their relative contributions to drug uptake at the subcellular level.

Our single-cell platform allows us to quantify heterogeneity in the cellular response to antibiotic treatment^45^. However, as detailed in the Results section, quantitative estimates of systematic and biological variation revealed no detectable heterogeneity in ofloxacin uptake in our experiments. Considering the large variations in gene and protein expression reported in bacterial cells and the corresponding heterogeneity in phenotypic traits including glucose uptake^19,35,46^, it is striking that ofloxacin uptake appears robust, i.e. uniform across cells within each of our experiments; however, a detailed investigation of this is beyond the scope of this study and will be further investigated in future work.

## Conclusions

We have developed a novel experimental and theoretical approach to study antibiotic accumulation label-free in individual Gram-negative bacteria in well-controlled microenvironments. Our experiments enabled us to quantify the role of the nutrient microenvironment and metabolic state of the cells in drug uptake at the single-cell level. We reported, to the best of our knowledge, the first quantitative comparisons between drug uptake in cells in different metabolic states and in cells with specific transport pathways disabled. Our experimental results showed that the growth phase of the cells, as determined by the nutrient microenvironment, plays a more significant role in ofloxacin uptake than either the porin OmpF or the efflux protein TolC. More generally, this suggests that the metabolic state of the cell is a crucial determinant of cellular drug uptake, which deserves detailed, quantitative investigation in well-controlled microenvironments. Combining our data with mathematical modelling and Bayesian inference enabled us to predict the kinetic parameters underlying ofloxacin accumulation in the different subcellular compartments of *E. coli* cells. This has previously proved extremely challenging primarily due to the small size of typical bacterial cells and the need for complicated washing steps before measuring drug uptake^12,13^, which may bias the results. We used the parameters extracted from fitting the model to our experimental data to predict drug accumulation in the outer membrane, the periplasm and the cytoplasm in parental, Δ*ompF* and Δ*tolC E. coli*.

Our approach offers possibilities for scaling up the number of drugs/pathogens that can be tested on the same chip, via parallelization of the cell trapping chambers. We also require small volumes of concentrated cultures for seeding the chip (<10 μl), which may facilitate its use in clinical settings. The assay also has the advantage of needing only micrograms of chemicals for testing, which is important when evaluating novel, candidate drugs that are typically expensive to manufacture. Our readout is based on fluorescence, and can be used to test the permeation properties of newly developed fluorescent antibiotic probes^47^, providing information about Gram-negative drug permeability for a range of different antibiotic classes. It could also be used to study the influence of specific functional groups on the uptake of closely related compounds. For instance, biophysical measurements of different fluoroquinolones revealed orders of magnitude differences in their lipid permeabilities^25^; our system facilitates similar studies on the bacteria themselves. The experimental setup is relatively simple to implement on standard epi-fluorescence microscopes and will provide researchers with a new, transferrable platform with which to study this vitally important permeation process in a range of pathogenic microbes.

## Materials and Methods

### Chemicals

Chemicals were purchased from Sigma-Aldrich unless otherwise stated. Ofloxacin stock solutions were prepared at a concentration of 10 mg/ml in 1 M NaOH. For the ofloxacin uptake experiments, the stock was diluted to a concentration of 12.5 μg/ml (100×MIC) in PBS. The minimal media used in the experiments was prepared in sterile water and contained 1×M9 salts, 2 mM MgSO_4_, 0.1 mM CaCl_2_ and 1 mg/L thiamine hydrochloride. The LB medium used for cell culture was the Melford high salt version containing 10 g/L casein digest peptone, 5 g/L yeast extract and 10 g/L NaCl; LB Agar plates were prepared with 15 g/L agar. Glucose stock solutions were prepared at a concentration of 0.5 M in sterile water and diluted to 1 g/L in minimal media for use in the experiments. Stock solutions of bovine serum albumin (BSA) were prepared at a concentration of 50 mg/ml in sterile water. A stock solution of propidium iodide (PI) was purchased from Thermo Fisher Scientific, and diluted 1:1000 in PBS for use in the experiments.

### Bacterial cell culture

All the *E. coli* strains used were BW25113 strains purchased from the Keio collection. The mutant strains contained kanamycin resistance cassettes in place of the deleted chromosomal gene. The strains were stored at −80 °C in a 1:1 ratio of overnight culture and 50% glycerol solution. 200 ml cultures were grown in LB (with 25 μg/ml kanamycin as necessary) at 37 °C overnight (with shaking at 200 rpm). Streak plates were prepared on LB agar (containing 25 μg/ml kanamycin as necessary), stored at 4 °C and used for a maximum of one week.

### Microfluidic chip fabrication

The complete protocol for the fabrication of the “mother-machine” microfluidic devices was reported previously^45^. The epoxy mold used was constructed from replicas of devices kindly provided by the Jun lab^48^. The final devices used were created by pouring polydimethylsiloxane (PDMS, Dow Corning, 9:1 base : curing agent) on to the epoxy mold; the PDMS was baked at 70 °C for 2 h in an oven. The PDMS chips were cut out and fluidic inlet/outlet columns punched using a 1.5 mm biopsy punch (Miltex). The PDMS chips were bonded to a type 1 coverslip using an air plasma treatment (10 s exposure at 30 W plasma power, Plasma etcher, Diener electronic GmbH, Germany) and left at 70 °C for 5 min to improve the adhesion. The chips were then filled with a 50 mg/ml solution of bovine serum albumin (BSA, in milliQ water) and incubated at 37 °C for 1 h. The BSA treatment passivates the internal surfaces of the chip thus preventing cells from adhering to the microchannels during experiments.

An overnight culture of cells (OD_595_ typically between 4.5-5) was resuspended in spent LB and concentrated to an OD of 50 (at 595 nm). The spent LB was prepared by centrifuging the overnight culture (10 min at 3000 g and 20 °C) – the supernatant was filtered twice through a 0.2 μm pore filter (Millipore). A 2 μl aliquot of this solution was injected into the microfluidic device and incubated at 37 °C for 20 min, enabling cells to enter the small side channels of the device. The filled device was then left overnight at room temperature before starting experiments.

### Drug uptake assay

Microfluidic flows were controlled using three parallel neMESYS syringe pumps (Cetoni GmbH, Germany) with glass syringes (ILS, Germany) of volumes 5 ml, 250 μl and 100 μl respectively. The syringes were interfaced with the microfluidic chips using FEP tubing (Upchurch Scientific 1520, I.D. = 0.03” and O.D. = 0.0625”). The syringes and the associated tubing were rinsed thoroughly with milliQ water and the appropriate experimental solutions before beginning the experiments, and with 70% ethanol after completion of the experiments.

All the experiments were performed on an Olympus IX73 epifluorescence microscope with an LED light source (wLS pE300, QImaging) using a 365 nm excitation wavelength LED. A standard DAPI filter set (Chroma ET series) modified with a ZET 365/20x excitation filter (Chroma) was used to better match the 365 nm excitation wavelength. An Olympus UPLSAPO 60×W (N.A 1.2) objective was used for all the experiments. We used a heating stage (Linkam Scientific THL60-16, UK) to maintain the cells at 37 °C throughout the experiments. All the ofloxacin experiments’ fluorescence intensity traces are presented in Figure S6 in the SI.

For the experiments on growing cells, chips containing initially non-growing *E. coli* were flushed with a continuous flow of fresh LB (100 μl/h) for 3 h, which led the cells to start growing and dividing. This was followed by a 10 min flush (at 300 μl/h) with minimal media containing 1 g/L glucose to wash away the LB. The glucose was added to the minimal media to prevent the cells from starving. Thereafter, ofloxacin (100×MIC, 12.5 μg/ml dissolved in PBS) was perfused through the chip at 100 μl/h, with images acquired at 5 s intervals using an Evolve 512 EMCCD camera (Photometrics) with 10 ms exposure times and an EM gain of 200 (bin 1, clearing mode – pre-exposure). The camera was controlled using μManager 1.4^49^. We chose to always dissolve the ofloxacin in PBS to ensure that the pH conditions remained uniform during drug exposure across all experiments and metabolic conditions; it is well known that pH regulates the charge state of fluoroquinolones, which affects their membrane permeabilities^25,50^. The LED was triggered by the camera to ensure that the cells were only exposed to the excitation light during image acquisition. It must be noted that to reduce the background auto-fluorescence at 365 nm, prior to the ofloxacin flush the imaging area was bleached with the excitation light for 5 s. As detailed below, we performed controls (see Figure S2) with propidium iodide staining after UV and ofloxacin exposure to confirm that the UV light used did not compromise the cells’ membrane integrity.

For experiments on non-growing cells, the chips containing non-growing *E. coli* were flushed for 10 min with PBS (300 μl/h) to wash away residual LB, the imaging area was bleached for 5 s with the UV light (365 nm) and subsequently the ofloxacin was perfused through the chip, with the drug concentration and imaging settings exactly the same as for the growing cell experiments.

For both growing and non-growing cell experiments, we performed auto-fluorescence controls where instead of the ofloxacin, PBS was perfused through the chip (the rest of the protocols remained identical). A representative dataset is reported in Figure S1(B) in the SI.

### Image Analysis

The image analysis was performed using a custom Python module^51^. First, a specified range of frames of the dataset are loaded. Optionally, manually selected out-of-focus time-points are ignored. Cell detection is performed on a frame-by-frame basis as follows. First the frame is filtered using a Difference-of-Gaussian (DoG) scale-space filter^52^ spanning a small range of scales, corresponding to the scale range of bacterial widths. The resulting scale-space volume is maximum-projected along the scale axis, and the automatic threshold detected using the Triangle method^53^.

The centroids of the regions in the binary image resulting from applying this threshold are used to determine the axis of the side channels by using Principal Component Analysis. The axis of the side channels is then used to determine the upper and lower extents of the side-channel-region, which are then used to generate a side-channel-region mask, in addition to two candidate main-channel-region masks. The side-channel-region mask is then used to select bacterial regions from the binary image. The correct channel is identified from the two candidate regions by analysing the fluorescence for the region whose mean signal exhibits the most variation.

Cells are tracked frame-to-frame by matching positions such that nearest-matching bacteria are assigned only if the match is cross-validated in both forward and backward temporal directions^54^. Bacterial trajectories are filtered to remove short trajectories (less than 10% of the full length).

The final trajectories are analysed as follows. First, a pre-determined dark-count (which is the average intensity of an image captured with the camera sensor covered) is subtracted from each bacterium’s mean fluorescence, yielding the dark-count-corrected mean intensities. The corresponding dark-count-corrected PDMS background values for each bacterium are obtained by averaging the pixel intensity values of the PDMS to the immediate left and right of the individual bacterium and applying a similar dark-count correction. This bacterium-specific dark-count-corrected PDMS background is subtracted from the corresponding bacterium. Finally, the background subtracted bacterium’s intensity at the starting time point is subtracted from all the values at later time points, yielding the background corrected bacterial fluorescence profiles over the course of the experiment (solid lines in Figures 2, S1 and S6).

For the drug dosage fluorescence, the initial intensity value of the dosage “main” channel (dark-count-corrected) is subtracted from all subsequent time points to initialise the drug fluorescence value to 0 (before drug arrival) – this also accounts for the subtraction of the background in the main channel. This reveals the drug dosage fluorescence profile over the course of the experiment (dashed lines in Figures 2, S1A,C and S6).

To account for any differences in absolute drug fluorescence between experiments, for the comparative analysis of drug uptake across the different experiments, all the background corrected cell and drug dosage fluorescence values in an experiment are normalised to the final value of the drug fluorescence in the main channel (*t* = 400 s) for that experiment. Note that this drug fluorescence value at *t* = 400 s is post-subtraction of the initial main channel background (measured before drug arrival) and thus always corresponds to the same concentration of ofloxacin (100×MIC, 12.5 μg/ml) across all experiments. These values are shown for a representative experiment in Figure 1D, and used for all comparative analysis (Figure 3) and modelling results in the paper. It is important to note that, since we are using this normalization in the model, we are assuming that the correspondence between drug fluorescence and concentration is the same in the main channel and in the vicinity of the cells. It is not possible to accurately resolve the drug fluorescence in the side channels in the immediate vicinity of each cell. The cells themselves are brighter than the surrounding channel and are hence easier to detect and track and, as specified above, we have established a protocol to subtract the scattering and fluorescence background for the cells.

Finally, since the cellular auto-fluorescence profiles were flat (Figure S1B,D), we did not need to correct for this effect when analysing the drug uptake experimental data; we simply subtracted the initial cellular fluorescence (at *t* = 0) from the cell fluorescence at all the time-points, as detailed above. We should also mention that the automated tracking works better for growing cells than for non-growing cells, which were smaller in size and therefore more difficult to detect. However, this does not significantly affect the average results, and the cell fluorescence values obtained through the automated code were similar to those obtained by manually selecting and measuring the cells in ImageJ; since we do not fit the model to the data for non-growing cells, we used the automated tracking results in all the figures in this manuscript.

### Quantifying intra-experimental variability

In order to estimate the variation in cellular fluorescence in the absence of the drug, we used the auto-fluorescence control experiment shown in Figure S1B to estimate the underlying biological and systematic variation in our experiments. These measurements report the *auto-fluorescence of the same cells* measured at different time points in the experiment. We quantified the coefficient of variation (CV) of the cell auto-fluorescence intensities (over the timescales of the experiment) of the 103 individual cells shown in Figure S1B. The *mean* CV across *all* the cells was 10 ± 3 % (*N* = 103, mean ± s.d.). This gives a quantitative estimate of the *measurement (systematic and underlying biological) heterogeneity for individual cells* within a single experiment.

We compare this variability in cellular auto-fluorescence with the apparent heterogeneity in drug uptake in the cells in Figure S1A. To estimate this value, we measured the intensity of the cells at the end of the drug uptake experiment (*t* = 400 s). The *heterogeneity in the cellular fluorescence corresponding to drug uptake* (in the knowledge that this includes the systematic and underlying biological variation mentioned above) is extracted by measuring the CV of the fluorescence across all the cells at this time-point. *Unlike* the CV measurement of the control which was for *individual cells* across *all* time-points, to estimate drug uptake heterogeneity amongst the 126 different cells, we measured the CV in the fluorescence of *all the cells* at the *final* time-point. This analysis yields a CV of 9.7%.

### Mathematical model

We model drug uptake in the different compartments of a Gram-negative bacterium (Figure 1A) using the following set of ordinary differential equations (ODEs):

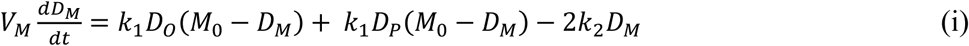

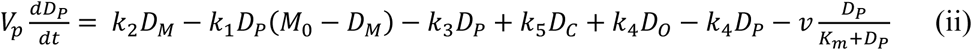

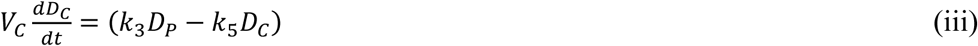

where *D*_*O*_, *D*_*M*_, *D*_*P*_ and *D*_*C*_ denote the drug concentrations in the external environment, the outer membrane, the periplasm and the cytoplasm, respectively. Importantly, we used the measured drug dosage traces for estimating *D*_*O*_ for every experiment, which allows us to control for any variations in the drug dosage profiles across different experiments (Figure S6). We model porin-mediated drug transport through the outer membrane as a two-step reversible process: drug molecules bind to porins with rate constant *k*_1_ from either side of the outer membrane and unbind to either side at rate *k*_2_. *M*_0_ denotes the concentration of functional porins in the outer membrane; based on literature values of the numbers of porins in typical Gram-negative outer membranes, we assumed that the total number of porins would vary between approximately 1×10^5^ to 2×10^5^ per cell (PS)^55^; this was used to restrict the range of possible values for *M*_0_. As a first approximation, we assume that diffusion through the LPS-lipid bilayer is negligible (*k*_4_∼0) in comparison to porin-mediated transport^12^. Furthermore, we postulate that ofloxacin molecules, like other fluoroquinolones^25,50^, diffuse across the inner membrane lipid bilayer (rate constants *k*_3_ and *k*_5_) and that the efflux of drug molecules from the periplasm to the external medium follows Michaelis-Menten kinetics with maximal rate *v* and Michaelis constant *K*_*m*_^31^. Parameters *V*_*M*_, *V*_*P*_ and *V*_*C*_ denote the volumes of the outer membrane, periplasm and cytoplasm, respectively (Table S2). The parameter *k*_3_ was calculated on the basis of passive diffusion measurements of ofloxacin permeability across lipid vesicle bilayers (Figure S3). To account for any potential binding of the drug to targets within the cytoplasm, we do not assume any equivalence between *k*_3_ and *k*_5_, an approach similar to that applied by Westfall *et al.*^31^; we only make the assumption that *k*_5_ ≤ *k*_3_. Crucially, the parameters (*k*_1_, *k*_2_, *k*_5_, *M*_0_, *K*_*m*_, *v*) were inferred from the experimental data obtained with the PS, Δ*ompF* and Δ*tolC E. coli* strains (Figure S6). The total drug concentration was calculated as:

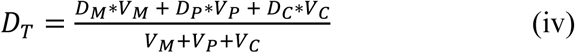

To model drug uptake in the Δ*ompF* strain, we used equations (i-iii) above, additionally assuming a possible decrease in the number of porins relative to the PS, i.e., *M*_0,Δ*ompF*_ ≤ *M*_0_. Similarly, for the case of the Δ*tolC* strain, we assumed that the *maximal* efflux rate may decrease relative to the PS, i.e., *v*_Δ*tolC*_ ≤ *v*.

All model simulations were run in Matlab (R2018b) using the in-built explicit Runge-Kutta (4, 5) solver (function ode45; default settings). The codes are available via GitHub.

### Parameter estimation

We obtained maximum likelihood estimates (MLEs) of the free model parameters (Table S2) using the medians of the drug uptake profiles for all the cells in an experiment. Please note that for convenience we use the term “population-averaged” throughout the text to refer to these *median* values of the drug uptake profiles. Since our data was normalized based on the fluorescence of the drug dose (see Methods; image analysis), estimates of parameters *k*_1_, *M*_0_, *K*_*m*_, *v* incorporate a constant factor related to the concentration of the drug dose (see Table S2). We denote the scaled version of these parameters using the prime symbol (′). We compiled a library of 18 datasets by combining population-averaged profiles from: (i) growing PS cells (3 experimental repeats); (ii) growing Δ*ompF* cells (3 experimental repeats) and (iii) growing Δ*tolC* cells (2 experimental repeats). We obtained parameter MLEs from each dataset, and to mitigate the risk of overfitting we then selected out of those parameter vectors the one that best fitted all 18 datasets. Under the assumption of Gaussian measurement error, the MLEs for each dataset correspond to parameter values minimising the following sum of squares: 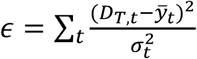. Here, 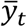 is the population-averaged drug uptake measurement at time *t*; *D*_*T,t*_ is the drug uptake predicted by the model; *σ*_*t*_ is the measurement error calculated based on a coefficient of variation of 4% (we obtained this from fluorescence measurements of the PDMS background); and the sum runs over all the time-points 0 to 400 s. Minimization was performed using Matlab’s in-built nonlinear least-squares solver (lsqcurvefit; with the maximum number of iterations set to 15). To find the global optimum of *ϵ*, we repeated the minimization task starting from 500 different initial points (generated using a Sobol sequence of quasi-random numbers) covering the entire parameter space.

We analyzed the single-cell data using a Bayesian hierarchical version of the model in which parameters *M*_0_ and *v* vary between single-cells. In particular, we postulate that these model-parameters are distributed at the population level according to two independent log-normal distributions^37^. Below, 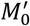 and *v*′ denote the rescaled versions of *M*_0_ and *v* which accommodate fitting the model to data normalized by the fluorescence of the drug dose (Table S2). The mean 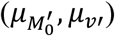 and standard deviation parameters 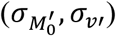 of each log-normal distribution dictate the average value of the corresponding model-parameter and its spread across a bacterial population. Posterior estimates of these population parameters 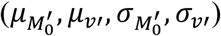 were inferred from single-cell data (experimental repeats were treated separately) using Gibbs sampling and informative priors based on the MLE estimates obtained in the step above (see Figures S4, S5 and Table S3 in the SI). In the first iteration (*j* = 1) of the algorithm, 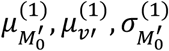, and 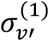 were drawn from their corresponding prior distributions and for each cell *i* = 1, …, *K* model-parameters 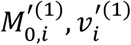 were obtained by minimizing the discrepancy between the model-predicted uptake profile and the single-cell measurements ***y***_*i*_ = {*y*_*i,t*_: *t* = 1, …, *Z*}. Subsequent iterations (*j* > 1) involve sampling in-turn from the full conditionals:

a. 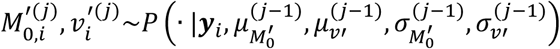;
b. 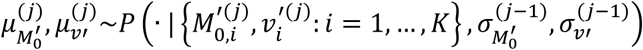;
c. 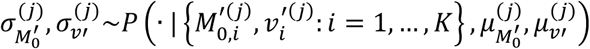.

In our analysis, we used conjugate priors for 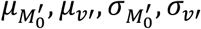, i.e., normal priors for 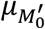 and *μ*_*v*′_, and gamma priors for 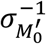 and 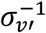. This choice greatly simplifies steps (b) and (c) as the target sampling distributions are the updated normal and gamma distributions, respectively. In step (a) for each cell *i* we sampled from the target distribution:

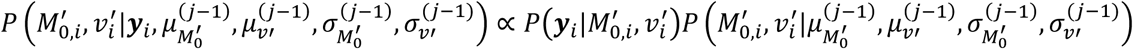

using a single Metropolis-Hasting step with a bivariate normal as the proposal distribution (covariance matrix set to 10^−4^**I**, where **I** is the 2×2 identity matrix). All results presented were obtained by running the Gibbs sampler for 2000 iterations (after having discarded 500 ‘warm-up’ iterations).

### Propidium Iodide (PI) staining to test membrane integrity after UV and ofloxacin treatment

To ensure that the combination of UV (365 nm) exposure and ofloxacin treatment does not compromise the cells’ membranes, we treated PS *E. coli* cells (growing) after an experiment with PI (1 μl dissolved in 1 ml PBS) for 10 min at a flow rate of 100 μl/h. PI is a stain commonly used to identify bacterial cells with compromised membranes. PI fluorescence was captured using an mCherry filter set (Chroma) using the green LED for excitation. A combined bright-field and mCherry fluorescence image representative of these experiments is shown in Figure S2, where it can be seen that less than 5% of the cells are stained with PI. Similar levels of PI staining were obtained for cells treated with ofloxacin but not bleached directly with the focused UV light. This suggests that our UV exposures do not compromise membrane integrity for the majority (>95%) of the cells.

## Author contributions

J.C. and M.V. contributed equally to this work. J.C. and S.P. conceptualized the project. J.C. performed all the experiments, with support from A.S. and J.I. The mathematical modelling and analysis was developed by M.V. and K.T.A. with inputs from J.C. and S.P.; the modelling was implemented by M.V. All the image analysis protocols and modules were developed by J.M. U.F.K. provided experimental resources and guidance. J.C., M.V. and S.P. wrote the manuscript, with input from all the other co-authors.

## Data and materials availability

All the data is available in the main text or the supplementary materials. The codes will be made available via a GitHub repository. Raw images may be made available to researchers upon reasonable request from the corresponding authors.

## References

1. Alberts, B. et al. Molecular Biology of the Cell. (Garland Science, 2008).

2. Missner, A. & Pohl, P. 110 years of the Meyer-Overton rule: Predicting membrane permeability of gases and other small compounds. Chemphyschem 10, 1405–1414 (2009).

3. Pages, J. M., James, C. E. & Winterhalter, M. The porin and the permeating antibiotic: a selective diffusion barrier in Gram-negative bacteria. Nat. Rev. Microbiol. 6, 893–903 (2008).

4. Zgurskaya, H. I., Rybenkov, V. V., Krishnamoorthy, G. & Leus, I. V. Trans-envelope multidrug efflux pumps of Gram-negative bacteria and their synergism with the outer membrane barrier. Res. Microbiol. 169, 351–356 (2018).

5. Liu, X. & Ferenci, T. Regulation of Porin-Mediated Outer Membrane Permeability by Nutrient Limitation in Escherichia coli. 180, 3917–3922 (1998).

6. Pu, Y. et al. Enhanced Efflux Activity Facilitates Drug Tolerance in Dormant Bacterial Cells. Mol. Cell 62, 284–294 (2016).

7. Gravelle, S. et al. Optimizing water permeability through the hourglass shape of aquaporins. Proc. Nati. Acad. Sci. USA 110, 16367–16372 (2013).

8. Berezhkovskii, A. M., Pustovoit, M. A. & Bezrukov, S. M. Channel-facilitated membrane transport: Transit probability and interaction with the channel. J. Chem. Phys. 116, 9952–9956 (2002).

9. Pagliara, S., Dettmer, S. L. & Keyser, U. F. Channel-facilitated diffusion boosted by particle binding at the channel entrance. Phys. Rev. Lett. 113, 048102 (2014).

10. Bleil, S., Reimann, P. & Bechinger, C. Directing Brownian motion by oscillating barriers. Phys. Rev. E 75, 031117 (2007).

11. Gladrow, J., Ritort, F. & Keyser, U. F. Experimental evidence of symmetry breaking of transition-path times. Nat. Commun. 10, 55 (2019).

12. Cama, J., Henney, A. M. & Winterhalter, M. Breaching the Barrier: Quantifying Antibiotic Permeability across Gram-Negative Bacterial Membranes. J. Mol. Biol. 431, 3531–3546 (2019).

13. Six, D. A., Krucker, T. & Leeds, J. A. Advances and challenges in bacterial compound accumulation assays for drug discovery. Curr. Opin. Chem. Biol. 44, 9–15 (2018).

14. O’Neill, J. Tackling drug-resistant infections globally: Final report and recommendations. (2016).

15. Silver, L. L. A Gestalt approach to Gram-negative entry. Bioorganic Med. Chem. 24, 6379–6389 (2016).

16. Mahendran, K. R. et al. Molecular basis of Enrofloxacin translocation through OmpF, an outer membrane channel of Escherichia coli - When binding does not imply translocation. J. Phys. Chem. B 114, 5170–5179 (2010).

17. Nestorovich, E. M., Danelon, C., Winterhalter, M. & Bezrukov, S. M. Designed to penetrate: time-resolved interaction of single antibiotic molecules with bacterial pores. Proc. Natl. Acad. Sci. U. S. A. 99, 9789–9794 (2002).

18. Du, D. et al. Structure of the AcrAB-TolC multidrug efflux pump. Nature 509, 512–515 (2014).

19. Ackermann, M. A functional perspective on phenotypic heterogeneity in microorganisms. Nat. Rev. Microbiol. 13, 497–508 (2015).

20. Vergalli, J. et al. Spectrofluorimetric quantification of antibiotic drug concentration in bacterial cells for the characterization of translocation across bacterial membranes. Nat. Protoc. 13, 1348–1361 (2018).

21. Piddock, L. J. V. & Johnson, M. M. Accumulation of 10 fluoroquinolones by wild-type or efflux mutant Streptococcus pneumoniae. Antimicrob. Agents Chemother. 46, 813–820 (2002).

22. Taheri-Araghi, S. et al. Cell-size Control and Homeostasis in Bacteria. Curr. Biol. 25, 385–391 (2015).

23. Hooper, D. C. Mechanisms of action and resistance of older and newer fluoroquinolones. Clin. Infect. Dis. 31, S24–S28 (2000).

24. Cama, J. et al. Quantification of Fluoroquinolone Uptake through the Outer Membrane Channel OmpF of Escherichia coli. J. Am. Chem. Soc. 137, 13836–13843 (2015).

25. Cama, J. et al. Direct Optofluidic Measurement of the Lipid Permeability of Fluoroquinolones. Sci. Rep. 6, 32824 (2016).

26. Sato, T. et al. Fluoroquinolone resistance mechanisms in an Escherichia coli isolate, HUE1, without quinolone resistance-determining region mutations. Front. Microbiol. 4, 125 (2013).

27. Zgurskaya, H. I., Krishnamoorthy, G., Ntreh, A. & Lu, S. Mechanism and function of the outer membrane channel TolC in multidrug resistance and physiology of enterobacteria. Front. Microbiol. 2, 189 (2011).

28. Widya, M. et al. Development and Optimization of a Higher-Throughput Bacterial Compound Accumulation Assay. ACS Infect. Dis. 5, 394–405 (2019).

29. Baba, T. et al. Construction of Escherichia coli K-12 in-frame, single-gene knockout mutants: the Keio collection. Mol. Syst. Biol. 2, 2006.0008 (2006).

30. Mahendran, K. R., Kreir, M., Weingart, H., Fertig, N. & Winterhalter, M. Permeation of antibiotics through Escherichia coli OmpF and OmpC porins: Screening for influx on a single-molecule level. J. Biomol. Screen. 15, 302–307 (2010).

31. Westfall, D. A. et al. Bifurcation kinetics of drug uptake by Gram-negative bacteria. PLoS One 12, e0184671 (2017).

32. Goldstein, R. E. Are theoretical results ‘Results’? Elife 7, e40018 (2018).

33. Asuquo, A. E. & Piddock, L. J. V. Accumulation and killing kinetics of fifteen quinolones for Escherichia coli, Staphylococcus aureus and Pseudomonas aeruginosa. J. Antimicrob. Chemother. 31, 865–880 (1993).

34. Richards, T. A., Massana, R., Pagliara, S. & Hall, N. Single cell ecology. Philos. Trans. R. Soc. B 374, 20190076 (2019).

35. Łapińska, U., Glover, G., Capilla-Lasheras, P., Young, A. J. & Pagliara, S. Bacterial ageing in the absence of external stressors. Philos. Trans. R. Soc. B 374, 20180442 (2019).

36. Hooper, D. C. Mechanisms of action of antimicrobials: focus on fluoroquinolones. Clin. Infect. Dis. 32, S9–S15 (2001).

37. Furusawa, C., Suzuki, T., Kashiwagi, A., Yomo, T. & Kaneko, K. Ubiquity of log-normal distributions in intra-cellular reaction dynamics. Biophysics (Oxf). 1, 25–31 (2005).

38. Kaščáková, S., Maigre, L., Chevalier, J., Réfrégiers, M. & Pagès, J. M. Antibiotic transport in resistant bacteria: Synchrotron UV fluorescence microscopy to determine antibiotic accumulation with single cell resolution. PLoS One 7, e38624 (2012).

39. Zampieri, M., Zimmermann, M., Claassen, M. & Sauer, U. Nontargeted Metabolomics Reveals the Multilevel Response to Antibiotic Perturbations. Cell Rep. 19, 1214–1228 (2017).

40. Smith, A. et al. The culture environment influences both gene regulation and phenotypic heterogeneity in Escherichia coli. Front. Microbiol. 9, 1739 (2018).

41. Keren, I., Kaldalu, N., Spoering, A., Wang, Y. & Lewis, K. Persister cells and tolerance to antimicrobials. FEMS Microbiol. Lett. 230, 13–18 (2004).

42. Reyes-Domínguez, Y., Contreras-Ferrat, G., Ramírez-Santos, J., Membrillo-Hernandez, J. & Carmen Gomez-Eichelmann, M. Plasmid DNA Supercoiling and Gyrase Activity in Escherichia coli Wild-Type and rpoS Stationary-Phase Cells. J. Bacteriol. 185, 1097–1100 (2003).

43. Sarathy, J., Dartois, V., Dick, T. & Gengenbacher, M. Reduced Drug Uptake in Phenotypically Resistant Nutrient-Starved Nonreplicating Mycobacterium tuberculosis. Antimicrob. Agents Chemother. 57, 1648–1653 (2013).

44. Swick, M. C., Morgan-Linnell, S. K., Carlson, K. M. & Zechiedrich, L. Expression of Multidrug Efflux Pump Genes acrAB-tolC, mdfA, and norE in Escherichia coli Clinical Isolates as a Function of Fluoroquinolone and Multidrug Resistance. Antimicrob. Agents Chemother. 55, 921–924 (2011).

45. Bamford, R. A. et al. Investigating the physiology of viable but non-culturable bacteria by microfluidics and time-lapse microscopy. BMC Biol. 15, 121 (2017).

46. Nikolic, N., Barner, T. & Ackermann, M. Analysis of fluorescent reporters indicates heterogeneity in glucose uptake and utilization in clonal bacterial populations. BMC Microbiol. 13, 258 (2013).

47. Stone, M. R. L., Butler, M. S., Phetsang, W., Cooper, M. A. & Blaskovich, M. A. T. Fluorescent Antibiotics: New Research Tools to Fight Antibiotic Resistance. Trends Biotechnol. 36, 523–536 (2018).

48. Wang, P. et al. Robust growth of Escherichia coli. Curr. Biol. 20, 1099–1103 (2010).

49. Stuurman, N., Amdodaj, N. & Vale, R. μManager: Open Source Software for Light Microscope Imaging. Micros. Today 15, 42–43 (2007).

50. Schaich, M. et al. An Integrated Microfluidic Platform for Quantifying Drug Permeation across Biomimetic Vesicle Membranes. Mol. Pharm. 16, 2494–2501 (2019).

51. Smith, A., Metz, J. & Pagliara, S. MMHelper: An automated framework for the analysis of microscopy images acquired with the mother machine. Sci. Rep. 9, 10123 (2019).

52. Lindeberg, T. Scale-space theory: A basic tool for analysing structures at different scales. J. Appl. Stat. 21, 225–270 (1994).

53. Zack, G. W., Rogers, W. E. & Latt, S. A. Automatic Measurement of Sister Chromatid Exchange Frequency. J. Histochem. Cytochem. 25, 741–753 (1977).

54. van der Walt, S. et al. scikit-image: image processing in Python. PeerJ 2, e453 (2014).

55. Delcour, A. H. Outer Membrane Permeability and Antibiotic Resistance. Biochim Biophys Acta. 1794, 808–816 (2009).

