## Supplemental Notes 1-2, Figures S1-S8, Tables S1-S4. for "Antibiotic transport kinetics in Gram-negative bacteria revealed via single-cell uptake analysis and mathematical modelling"

**This PDF file includes:**

Supplemental Notes 1-2
Figures S1 to S8
Tables S1 to S4

### Supplemental Note 1: Quantifying drug dosage

Our experimental approach, using syringe pumps to control drug delivery and auto-fluorescence for drug detection (Methods), enables us to precisely quantify the arrival of the drug in the vicinity of the cells under investigation. Representative images of the main, drug delivery channel and the side, cell-hosting channels before and after drug dosage are shown in Figure 1C (main text). Example drug dosage profiles are reported in Figure 2 of the main text (dashed black lines) and in the SI (dashed black lines in Figures S1A,C and S6). Since the drug dosage concentration is the same across all experiments, this measurement allows us to correct for any drug fluorescence intensity variation across all the different experiments (Methods). We measure the background at $t = 0$, and observe that the drug arrives in the field of view typically around 100 s after the start of the experiment, reaching its final concentration around 200 s (Figure 2A-D). We use the final, steady-state value of the drug dose fluorescence (at $t = 400$ s) to map drug fluorescence to drug concentration (see Methods). Importantly, since we measure the drug dosage profile across different experiments, we use this information as an input to the model, which allows us to account for any differences between the dose profiles across the different experiments.

### Supplemental Note 2: Quantifying cellular autofluorescence

For each strain/condition, we performed control experiments to measure the auto-fluorescence profiles of individual bacteria in the absence of the drug (see Methods). A representative comparison between cellular drug fluorescence and auto-fluorescence profiles is reported in Figure S1, corresponding to either the presence (Figure S1A,C) or absence (Figure S1B,D) of the drug. We observe that the cellular control auto-fluorescence profiles are flat across the timescales of the experiment; thus cellular auto-fluorescence has a negligible effect on the drug uptake profiles. Similar cellular auto-fluorescence profiles were observed across all the control experiments performed (number of experimental repeats: 2 (PS growing), 3 (Δ*ompF* growing), 2 (Δ*tolC* growing), 3 (PS non-growing) and 3 (Δ*tolC* non-growing)).

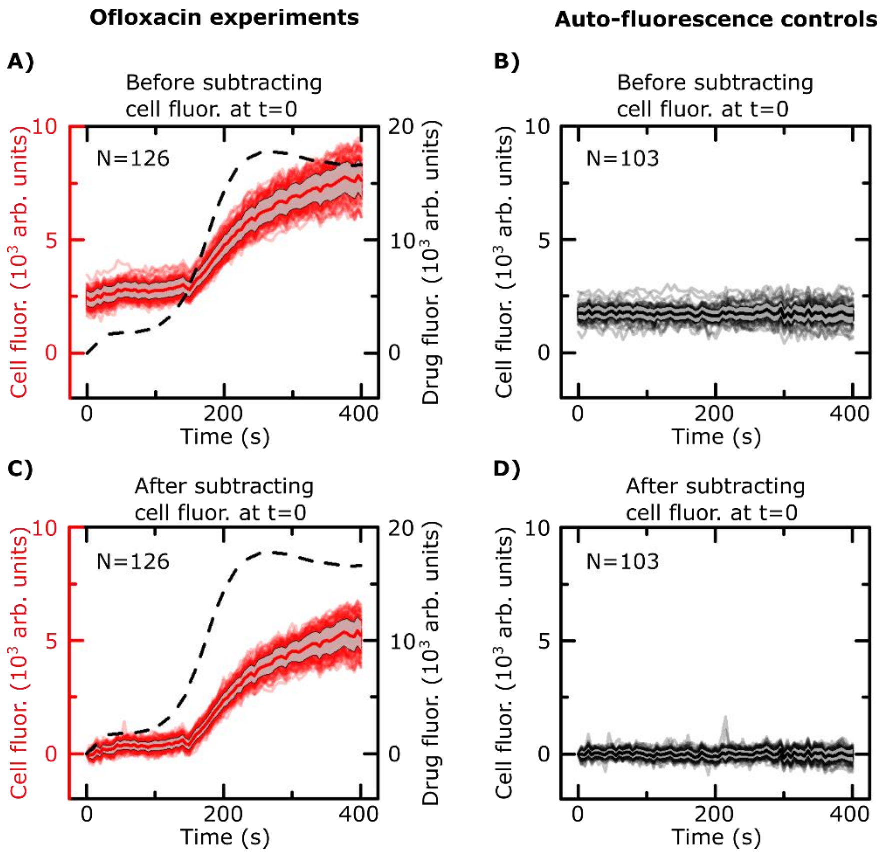


**Fig. S1.**

**Comparison of drug fluorescence (A, C) versus auto-fluorescence control (B, D) cellular profiles of parental strain (PS) *E. coli* (growing).** In the top panels (A and B), the cellular fluorescence profiles report the fluorescence intensities of individual bacteria after the subtraction of the PDMS background fluorescence as explained in the Methods. The bottom two panels (C) and (D) report the fluorescence intensities of the cells in the same experiments as reported in (A) and (B), respectively, after *also* subtracting the initial cell fluorescence values at $t = 0$. In both experiments, the cells were grown on chip for 3 h in fresh LB medium at a flow rate of 100 μl/h, followed by a wash (10 min at 300 μl/h) with 1 g/L glucose dissolved in minimal media prior to dosage with ofloxacin (A, C) or PBS (B, D). The drug fluorescence experiment (A, C) shows the delivery (dashed black line) of ofloxacin in the main channel and corresponding drug uptake profile of 126 PS *E. coli* cells (red); the mean (thick red line) and standard deviation (grey shading) for the cell profiles are also shown. In the absence of the drug, the cellular auto-fluorescence profiles are flat (B, D; 103 individual cells shown in grey with the mean in black, along with the standard deviation as the grey shaded region). We conclude that cellular auto-fluorescence has a negligible contribution to the drug fluorescence profile of a cell under the conditions of our experiment.

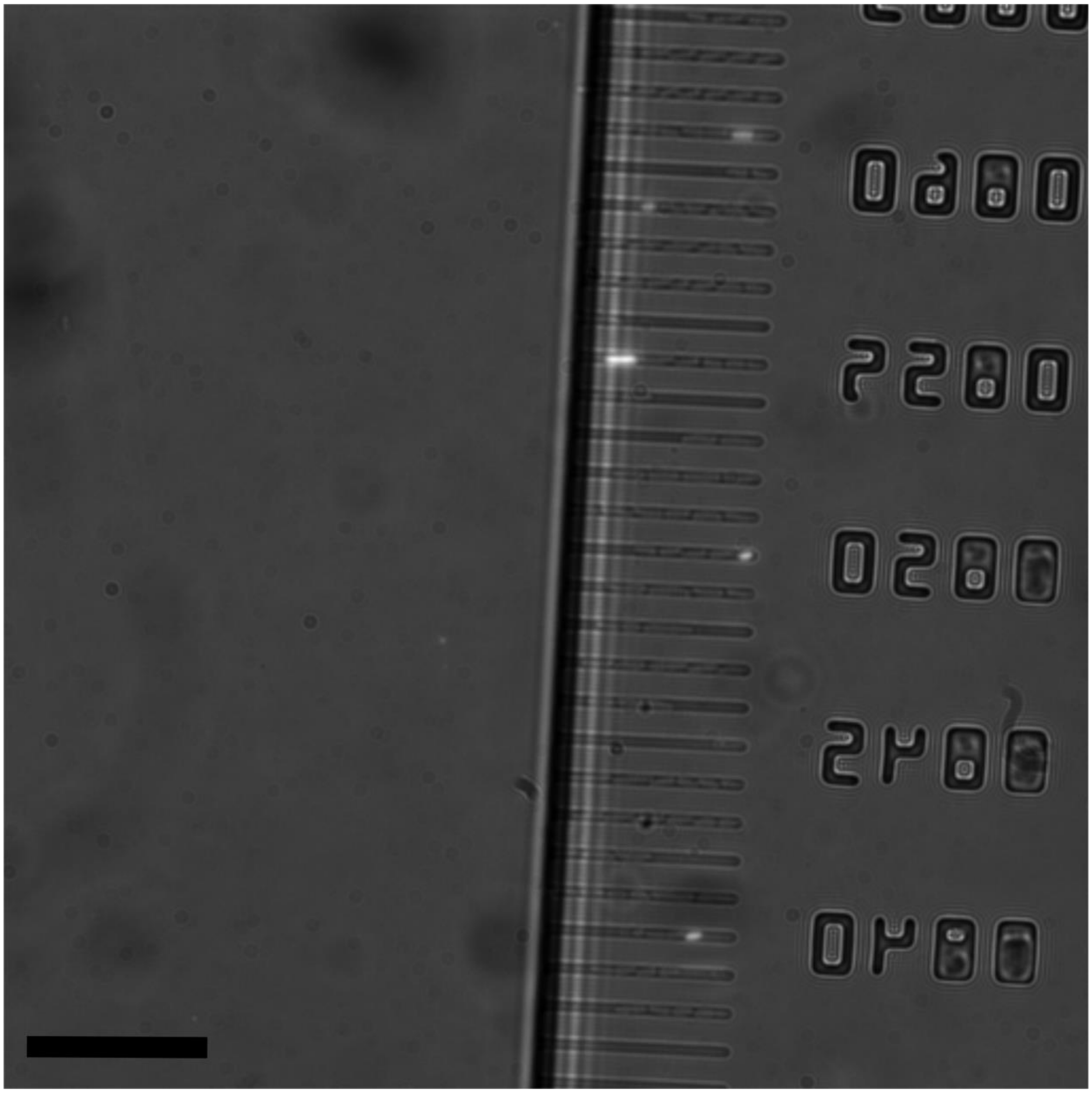

**Fig. S2.**

**Representative combined bright-field and fluorescence image showing the effects of Propidium Iodide (PI) staining on the bacteria post UV and ofloxacin exposure.** The PI test shows that cellular membrane integrity remains intact during the drug uptake experiments. PS *E. coli* cells were grown for 3 h on fresh LB in the chip and treated with UV ($\lambda_{ex}$ = 365 nm) and ofloxacin as per the standard drug uptake experimental protocol (Methods). After the ofloxacin treatment, the cells were treated with PI for 10 min (flow rate 100 μl/h). Less than 5% of the cells stain with PI; similar levels of PI staining were obtained in cells that did not receive the focused UV treatment. Our results indicate that the UV exposure ($\lambda_{ex}$ = 365 nm) does not damage cellular membrane integrity for the majority (>95%) of the cells within the timescale of the drug uptake experiments. Scale bar = 25 μm.

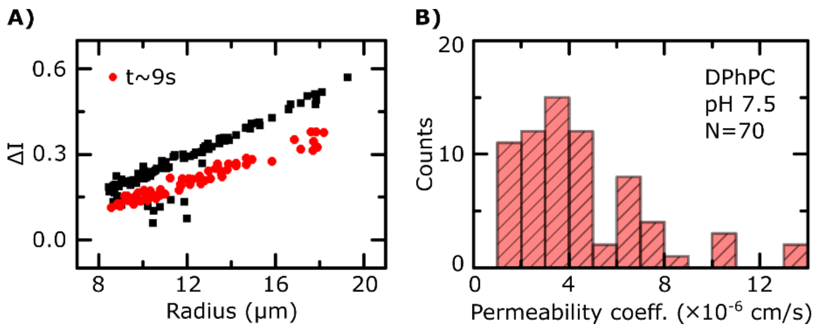


**Fig. S3.**

**Ofloxacin permeability measurement across DPhPC lipid vesicle membranes at pH 7.5.** Lipid vesicles were prepared by electroformation and treated with a solution of 2 mM ofloxacin in a T-junction microfluidic chip (for technical details of the measurement conditions and assay, please refer to Cama *et al. Sci. Reps.* 2016). Drug permeation across the vesicle membranes is tracked using the UV auto-fluorescence of ofloxacin. **A)** $\Delta I$ refers to a normalized intensity difference between the interior and exterior of the vesicles. The shift in $\Delta I$ between vesicles detected at $t$ = 0 (black squares) and those detected at a later time point ($t$ ~ 9 s, red circles) is a direct readout of drug transport into the vesicles, associated with an increase in the fluorescence intensity of the vesicles due to the accumulation of the drug. **B)** Permeability coefficient histograms associated with the measurement of ofloxacin uptake in 70 vesicles. The permeability coefficient of ofloxacin is calculated based on a simple diffusion model (details in Cama *et al. Sci. Reps. 2016*, Cama *et al. Lab Chip* 2014). The permeability coefficient measured was $P = 4.5 \pm 0.3 \times 10^{-6}$ cm/s ($N$ = 70, mean ± s.e.m.).

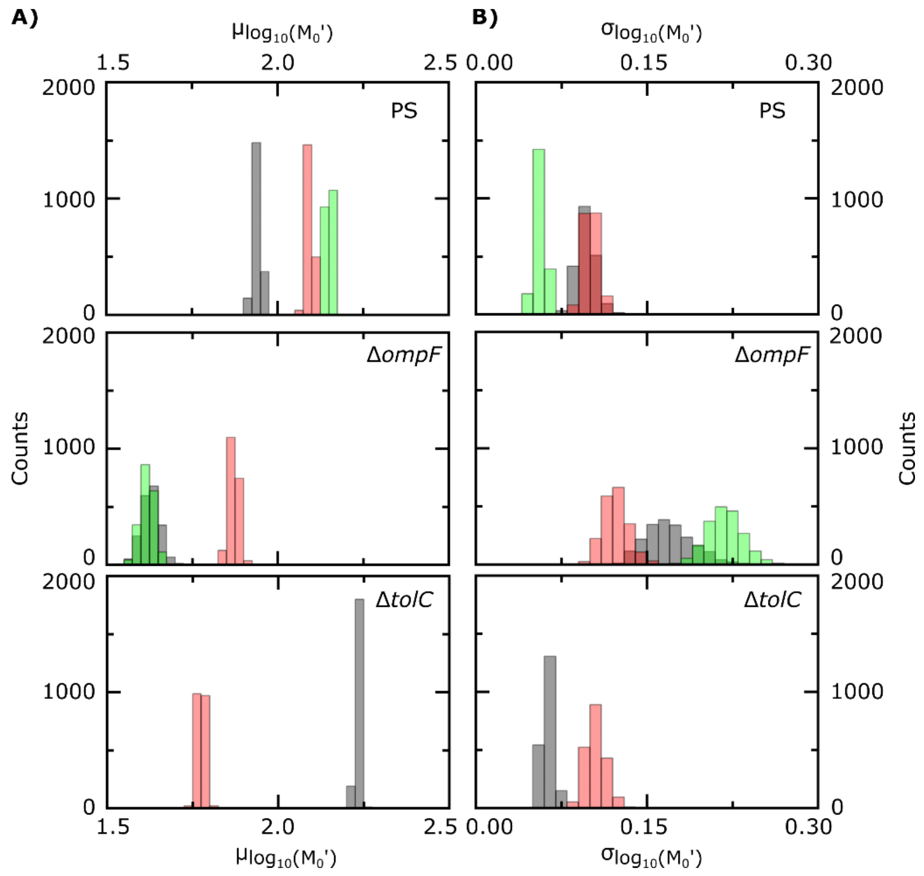


**Fig. S4.**

**Posterior distributions for the means (A) and standard deviations (B) of the log-normal distributions for $M_0'$ in individual experiments.** Individual experimental repeats are represented in different colors for each *E. coli* strain.

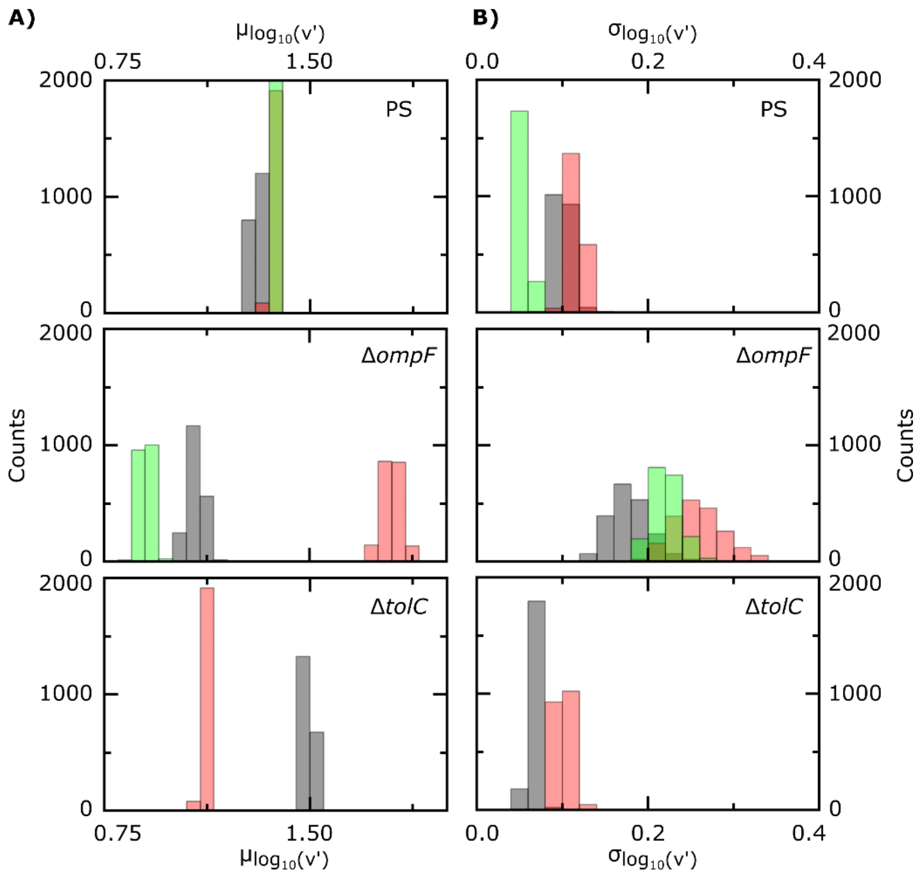


**Fig. S5.**

**Posterior distributions for the means (A) and standard deviations (B) of the log-normal distributions for $v'$ in individual experiments.** Individual experimental repeats are represented in different colors for each *E. coli* strain.

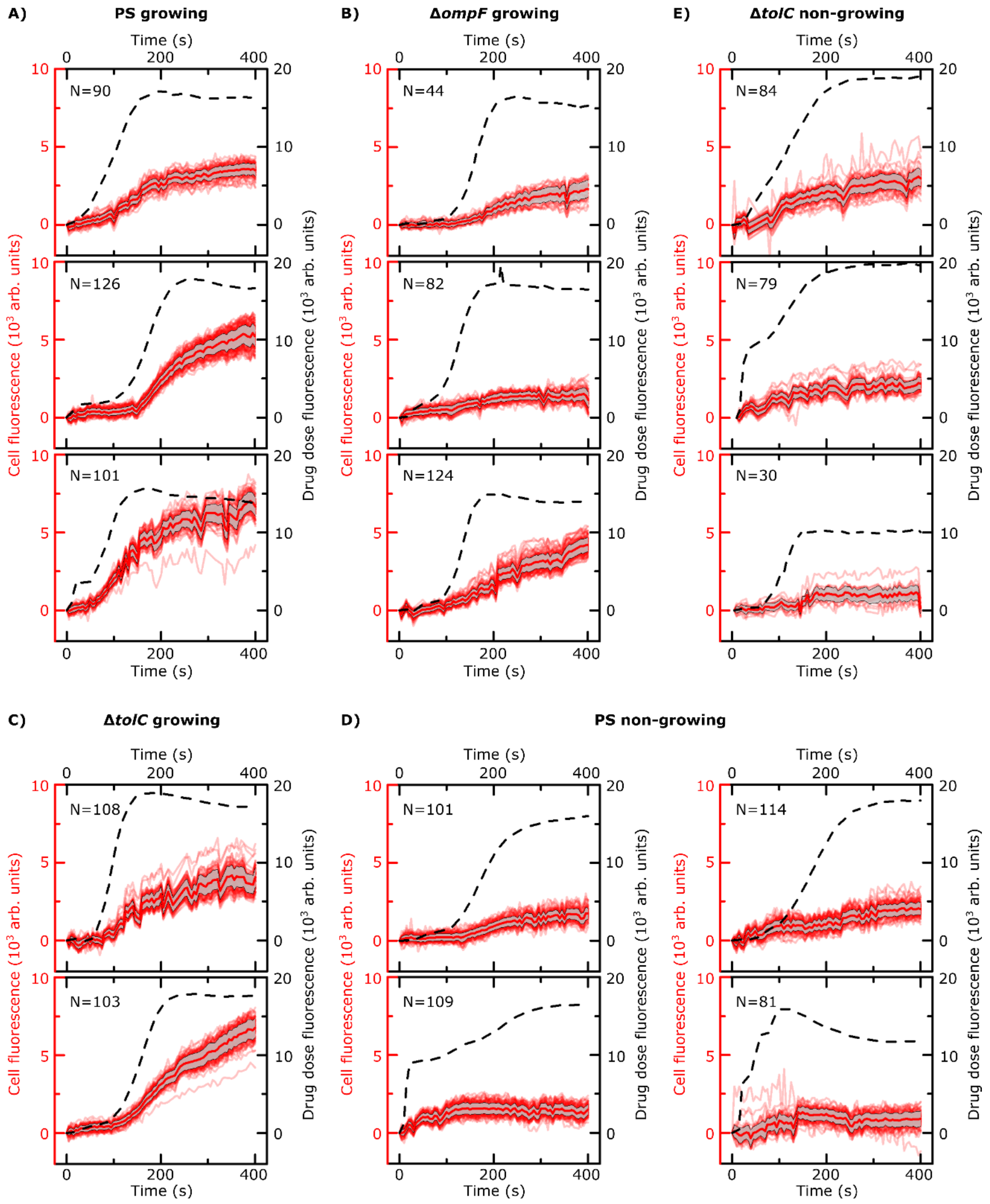


**Fig. S6.**

**Complete dataset showing all experimental repeats for the ofloxacin uptake experiments.** Each plot corresponds to an individual repeat, with the strains/conditions distributed in panels (A)-(E). Black dashed lines correspond to drug dosage profiles (right Y-axes), red lines correspond to cellular fluorescence profiles (left Y-axes) with the mean (thick red line) and standard deviation (grey shading) also shown in the individual plots. *N* refers to the number of cells in the individual experiment. All values reported after subtracting the corresponding backgrounds and the cell fluorescence at $t = 0$ (see Methods).

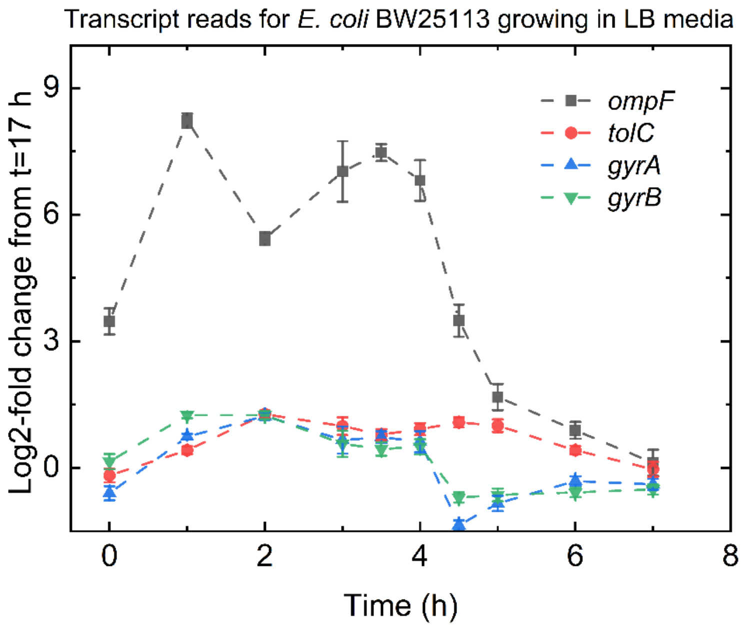


**Fig. S7.**

**Population level transcriptomic data for the *ompF*, *tolC*, *gyrA* and *gyrB* genes of *E. coli* BW25113 (parental strain) grown in LB media (10g/L NaCl).** Reproduced using data made available in Smith *et al.*, *Front. Microbiol.* 2018. The time axis refers to the time of growth in LB following the seeding of a stationary phase (17 h of growth in LB) culture into fresh LB. Dashed lines connecting the data points are provided as guides for the eye. Error bars represent the standard error of the mean.

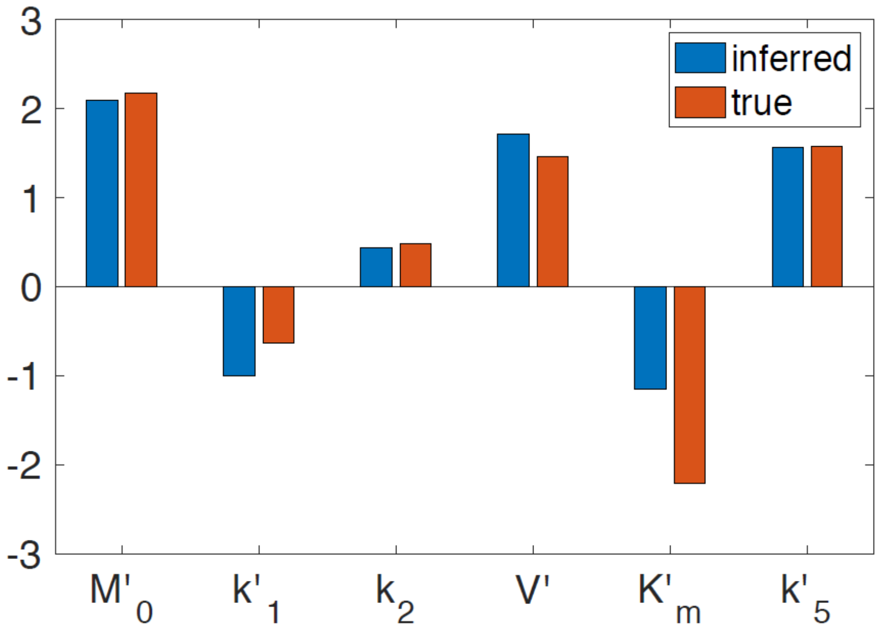


**Fig. S8.**

**Comparison between parameter log10-values inferred from simulated data (inferred, blue) and the actual values used to simulate the data (true, red).** We used the model to simulate single-cell drug uptake profiles similar to those we observed experimentally for PS *E. coli*, and the Δ*ompF* and Δ*tolC* mutant strains. The parameter values we used can be found in Table S2. To introduce cell-to-cell heterogeneity, we assumed that parameters $M_0'$ and $v'$ varied within the different populations according to the distributions found in Figures 4A and 4B. Parameter values were recovered from population-averaged profiles using the methodology described in the Methods.

| Comparison of whole cell fluorescence (normalized) at $t$ = 400 s | p-value (2 sample t-test with Welch's correction) |
|---|---|
| PS non-growing vs PS growing | $1.4 \times 10^{-123}$ |
| PS non-growing vs Δ*ompF* growing | $1.2 \times 10^{-33}$ |
| PS non-growing vs Δ*tolC* growing | $1.3 \times 10^{-98}$ |
| PS non-growing vs Δ*tolC* non-growing | $1.9 \times 10^{-5}$ |
| PS growing vs Δ*ompF* growing | $1.8 \times 10^{-42}$ |
| PS growing vs Δ*tolC* growing | $2.7 \times 10^{-4}$ |
| PS growing vs Δ*tolC* non-growing | $8.5 \times 10^{-112}$ |
| Δ*ompF* growing vs Δ*tolC* growing | $7.4 \times 10^{-30}$ |
| Δ*ompF* growing vs Δ*tolC* non-growing | $2.5 \times 10^{-21}$ |
| Δ*tolC* growing vs Δ*tolC* non-growing | $3.2 \times 10^{-92}$ |

**Table S1.**

**Two sample t-test (with Welch's correction) p-values for the whole cell fluorescence (background subtracted, normalized) of the different strains/conditions at *t* = 400 s (Figure 3B). Sample sizes: PS non-growing (*N* = 405), PS growing (*N* =317), Δ*ompF* growing (*N* = 250), Δ*tolC* growing (*N* = 211) and Δ*tolC* non-growing (*N* = 193).**

| Parameter | Units | Value | Notes |
|---|---|---|---|
| $k_1$ | $\text{molecules}^{-1} \cdot \mu m^6 \cdot \sec^{-1}$ | $1.12\times10^{-5}$ | Inferred from cell data. |
| $k_2$ | $\mu\text{m}^3 \cdot \sec^{-1}$ | 3.08 | Inferred from cell data. |
| $k_3$ | $\mu\text{m}^3 \cdot \sec^{-1}$ | 0.52 | Estimated from data in Fig. S3. |
| $k_4$ | $\mu\text{m}^3 \cdot \sec^{-1}$ | 0 | Assumed negligible. |
| $k_5$ | $\mu\text{m}^3 \cdot \sec^{-1}$ | 0.0139 | Inferred from cell data. |
| $M_0$ | $\text{molecules} \cdot \mu m^{-3}$ | $3.11\times10^6$ | Inferred from cell data. |
| $v$ | $\text{molecules} \cdot \sec^{-1}$ | $6.11\times10^5$ | Inferred from cell data. |
| $K_m$ | $\text{molecules} \cdot \mu\text{m}^{-3}$ | $1.30\times10^2$ | Inferred from cell data. |
| $V_M$ | $\mu\text{m}^3$ | 0.0565 | The bacterial cell is modelled as a cylinder of total width 1 μm. The width of the outer membrane was estimated to be 5 nm, and that of the periplasmic space 30 nm. |
| $V_P$ | $\mu\text{m}^3$ | 0.326 | |
| $V_C$ | $\mu\text{m}^3$ | 2.45 | |
| Model fitting | | | |
| Parameters | $k_1' = k_1 \cdot A\,; M_0' = \frac{M_0}{A}; v' = \frac{v}{A}; K_m' = \frac{K_m}{A};$ $k_2; k_5' = k_3/k_5$ | | Parameterization for fitting the model with fluorescence data. $A = 2.08 \times 10^4$ molecules $\cdot \mu m^{-3}$ is the drug dose concentration. |
| For the estimation of parameters from the population-averaged data, to ease exploration of the parameter space, all parameters above were log (base 10) transformed. The range of parameter $\log_{10}k_5'$ was set to [0,5] to satisfy the constraint $k_5 \leq k_3$. The range of parameter $M_0$ was constrained in [1.93, 2.23] so that the corresponding count of porins per cell lies between $1\times10^5$ to $2\times10^5$. All other parameters we allowed to vary between [-5, 5]. | | | |

**Table S2.**

**Model parameter estimates for parental strain *E. coli*.**

| Parameter | Values |
| --- | --- |
| $\boldsymbol{\mu}_{\log_{10}(M_0')}$ | $\boldsymbol{\mathcal{N}}(\mu = 2.175, \sigma = 0.5)$ |
| $\boldsymbol{\mu}_{\mathbf{log_{10}}(v')}$ | $\boldsymbol{\mathcal{N}}(\mu = 1.47, \sigma = 0.5)$ |
| $\boldsymbol{\sigma}_{\log_{10}(M_0')}$ | $\Gamma(\alpha = 10^{-4}, \beta = 10^{-4})$ |
| $\boldsymbol{\sigma}_{\mathbf{log_{10}}(v')}$ | $\Gamma(\alpha = 10^{-4}, \beta = 10^{-4})$ |

**Table S3.**

**Prior distributions for population parameters in the Bayesian hierarchical model.** Normal distributions with mean $\boldsymbol{\mu}$ and st. dev. $\boldsymbol{\sigma}$ were used as priors for $\boldsymbol{\mu}_{\log_{10}(M_0')}$ and $\boldsymbol{\mu}_{\mathbf{log_{10}}(v')}$. Gamma distributions with shape parameter $\boldsymbol{\alpha}$ and scale parameter $\boldsymbol{\beta}$ were used as priors for $\boldsymbol{\sigma}_{\log_{10}(M_0')}$ and $\boldsymbol{\sigma}_{\mathbf{log_{10}}(v')}$.

| Probability | Value |
| --- | --- |
| $D_M^{\mathrm{PS}} > D_M^{\Delta ompF}$ | 0.924 |
| $D_M^{\mathrm{PS}} > D_M^{\Delta tolC}$ | 0.525 |
| $D_P^{\mathrm{PS}} > D_P^{\Delta ompF}$ | 0.718 |
| $D_P^{\mathrm{PS}} > D_P^{\Delta tolC}$ | 0.549 |
| $D_C^{\mathrm{PS}} > D_C^{\Delta ompF}$ | 0.719 |
| $D_C^{\mathrm{PS}} > D_C^{\Delta tolC}$ | 0.549 |
| $D_T^{\mathrm{PS}} > D_T^{\Delta ompF}$ | 0.748 |
| $D_T^{\mathrm{PS}} > D_T^{\Delta tolC}$ | 0.555 |

**Table S4.**

**Using the model to compare drug accumulation in different strains after 400 s of drug exposure.** The model was used to estimate the probability of higher drug accumulation within PS cells compared to Δ*tolC* and Δ*ompF* mutant cells at the subcellular and whole cell level (M, outer membrane; P, periplasm; C, cytoplasm; T, total). Probability estimates are based on 1000 runs of the model for each strain in which parameters $M_0'$ and $v'$ were drawn from the corresponding population distributions shown in Figure 4 of the main text.
